# Deep learning-guided selection of antibody therapies with enhanced resistance to current and prospective SARS-CoV-2 Omicron variants

**DOI:** 10.1101/2023.10.09.561492

**Authors:** Lester Frei, Beichen Gao, Jiami Han, Joseph M. Taft, Edward B. Irvine, Cédric R. Weber, Rachita K. Kumar, Benedikt N. Eisinger, Sai T. Reddy

**Affiliations:** Department of Biosystems Science and Engineering, ETH Zurich; Basel 4058, Switzerland; Botnar Research Centre for Child Health; Basel 4058, Switzerland; Alloy Therapeutics (Switzerland) AG, Allschwil 4123, Switzerland

**Author notes:** equal contribution (Lester Frei, Beichen Gao).

## Abstract

Most COVID-19 antibody therapies rely on binding the SARS-CoV-2 receptor binding domain (RBD). However, heavily mutated variants such as Omicron and its sublineages, which are characterized by an ever increasing number of mutations in the RBD, have rendered prior antibody therapies ineffective, leaving no clinically approved antibody treatments for SARS-CoV-2. Therefore, the capacity of therapeutic antibody candidates to bind and neutralize current and prospective SARS-CoV-2 variants is a critical factor for drug development. Here, we present a deep learning-guided approach to identify antibodies with enhanced resistance to SARS-CoV-2 evolution. We apply deep mutational learning (DML), a machine learning-guided protein engineering method to interrogate a massive sequence space of combinatorial RBD mutations and predict their impact on angiotensin-converting enzyme 2 (ACE2) binding and antibody escape. A high mutational distance library was constructed based on the full-length RBD of Omicron BA.1, which was experimentally screened for binding to the ACE2 receptor or neutralizing antibodies, followed by deep sequencing. The resulting data was used to train ensemble deep learning models that could accurately predict binding or escape for a panel of therapeutic antibody candidates targeting diverse RBD epitopes. Furthermore, antibody breadth was assessed by predicting binding or escape to synthetic lineages that represent millions of sequences generated using *in silico* evolution, revealing combinations with complementary and enhanced resistance to viral evolution. This deep learning approach may enable the design of next-generation antibody therapies that remain effective against future SARS-CoV-2 variants.

## INTRODUCTION

The onset of the COVID-19 pandemic spurred the rapid discovery, development and clinical approval of several antibody therapies. The monoclonal antibody LY-CoV555 (bamlanavimab) (Eli Lilly)^1^ and the combination therapy consisting of REGN10933 (casirivimab) and REGN10987 (imdevimab) (Regeneron)^2^ were among the first to receive Emergency Use Authorization (EUA) from the United States FDA in late 2020. The primary mechanism of action for these therapies consist of virus neutralization by binding to specific epitopes of the RBD of SARS-CoV-2 spike (S) protein, thus inhibiting viral entry into host cells via the ACE2 receptor. However, the emergence of SARS-CoV-2 variants such as Beta, Gamma and Delta, each characterized by numerous mutations in the RBD, exhibited reduced sensitivity to neutralizing antibodies, including LY-CoV555^3,4^, whose EUA was subsequently revoked. Of note, antibody combination therapies such as those from Regeneron and Eli Lilly (LY-CoV555+LY-CoV16 (etesevimab)) were more resilient to viral variants and maintained their EUA throughout most of 2021^3^. However, the emergence and rapid spread of Omicron BA.1 in late 2021, a variant which has a staggering 35 mutations in the S protein, 15 of which are in the RBD resulted in substantial escape from nearly all clinically approved antibody therapies^5^. This includes the combination therapies from Regeneron and Eli Lilly, which also had their EUAs subsequently revoked^6^. Even antibody therapies with exceptional breadth, which were initially discovered against the ancestral SARS-CoV-2 (Wu-Hu-1) and retained neutralizing activity against BA.1 – S309 (sotrovimab) (GSK/Vir)^7^ and LY-CoV1404 (bebtelovimab) (Eli Lilly)^8^ – lost efficacy against subsequent Omicron sublineages (e.g., BA.2, BA.4/5, and BQ.1.1)^9,10^ and had their clinical use authorization revoked. Despite there being a critical need for antibody therapies for the protection of at-risk populations (young children, the elderly, individuals with chronic illnesses, and those with weakened immune systems)^11–15^, since March 2023, there are no antibody therapies with an active clinical authorization for COVID-19^16^.

The ephemeral clinical life span of COVID-19 antibody therapies has emphasized that, in addition to established metrics for antibody therapeutics (e.g. neutralization potency, affinity, and developability)^17^, it is imperative to evaluate antibody breadth (ability of an antibody to bind to divergent SARS-CoV-2 variants) at early stages of clinical development. This may enable selection of lead candidates that have the most potential to maintain activity against a rapidly mutating SARS-CoV-2. To address this, high-throughput protein engineering techniques such as deep mutational scanning (DMS)^18^ have been extensively employed to profile the impact of single position mutations in the RBD on ACE2-binding and antibody escape^5,19–24^. While DMS has proven effective for profiling single mutations, many SARS-CoV-2 variants that have emerged possess multiple mutations in the RBD. For example the aforementioned Omicron BA.1 lineage, or the recently identified BA.2.86, which possesses an astonishing 13 RBD mutations relative to its closest Omicron variant (BA.2) and 26 RBD mutations relative to ancestral Wu-Hu-1^25–27^. Experimental screening of combinatorial RBD mutagenesis libraries (e.g., using yeast surface display) vastly undersamples the theoretical protein sequence space, therefore computational approaches are increasingly being employed in concert. For instance, experimental measurements such as DMS data have been used to calculate statistical estimators^28^ or to train machine learning models that make predictions on ACE2 binding and antibody escape^29–31^. While such computational tools enable interrogation of a larger mutational landscape of SARS-CoV-2, their primary reliance on datasets that largely consist of single mutations from DMS experiments limits their ability to capture the effects of combinatorial mutations, especially in the context of high mutational variants such as Omicron sublineages (e.g., BA.1, BA.4/5, BA.2.86).

Here, we apply deep mutational learning (DML), which combines yeast display screening, deep sequencing and machine learning to address the emergence of Omicron BA.1 and its many sublineages. We expand the scope of DML from screening short, focused mutagenesis libraries^32^ to screening combinatorial libraries spanning the entire RBD for binding to ACE2 or binding/escape from antibodies. Ensemble deep learning models utilizing dilated residual network blocks were trained with deep sequencing data and shown to make accurate predictions for ACE2 binding and antibody escape. Next, deep learning was used to determine the breadth of second-generation antibodies (with known binding to BA.1) across a massive sequence landscape of BA.1-derived synthetic lineages, allowing the rational selection of specific antibody combinations that optimally cover the RBD mutational sequence space. This approach provides a powerful tool to guide the selection of antibody therapies that have enhanced resistance to both current and future high mutational variants of SARS-CoV-2.

## RESULTS

### Design and construction of a high distance Omicron BA.1 RBD library

A mutagenesis library was constructed based on BA.1, covering the entire 201 amino acid (aa) RBD region (positions 331 - 531 of SARS-CoV-2 S protein). To maximize the interrogated RBD sequence space, the library design was entirely synthetic and unbiased, as it did not consider evolutionary data or previous experimental findings. For the construction of the library, the RBD sequence was split into 11-12 fragments, each with an approximate length of 48 nucleotides (nt) (Supplementary Table 1). For each fragment, 136 different single-stranded oligonucleotides (ssODN) were designed, where each ssODN had either one codon or all combinations of two codons replaced by fully degenerate NNK codons (N = A, G, C, or T; K = G or T) (Fig. 1a) (Methods). For each fragment, ssODNs were amplified using PCR to generate double-stranded DNA. Each fragment was flanked by recognition sites for the type II-S restriction enzyme BsmBI, thus enabling assembly into full-length RBD regions by Golden Gate assembly (GGA)^33^. GGA utilizes type II-S restriction enzymes capable of cleaving DNA outside their recognition sequence, thereby allowing the resulting DNA overhangs to have any sequence. Based on the overhangs, individual fragments were assembled by DNA ligase to full-length RBD sequences with high fidelity^34,35^. The restriction sites were eliminated during the process, thus enabling scarless assembly of full length RBD sequences (Fig. 1b, Methods)^34^. This approach yielded approximately 98% correctly assembled RBD sequences (Supplementary Fig. 1). Since GGA required four nt homology between individual fragments for ligation, this led to portions of the sequence which needed to remain constant, thereby restricting library diversity^36^. To overcome this limitation, four sub-libraries were designed and individually assembled. Using sub-library 1 as a reference, sub-library 2 is shifted by 12 nt, sub-library 3 by 24 nt and sub-library 4 by 36 nt. These sub-libraries provided an increase in the mutational space covered by the RBD combinatorial mutagenesis library, since at the GGA homology for a given library, the remaining three libraries can have mutations (Fig. 1c).

**Figure 1.**
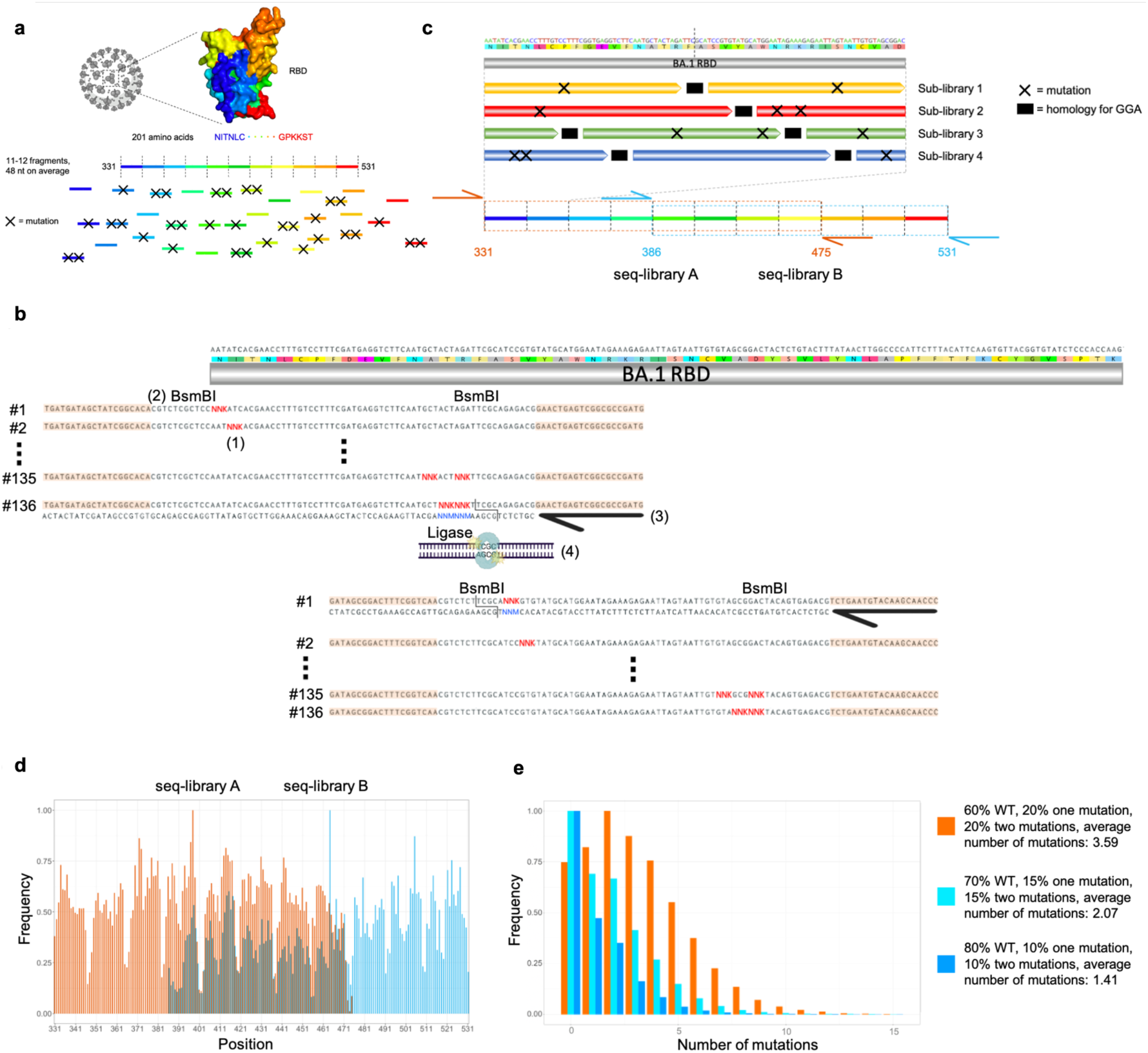
Construction of a high edit distance (ED) synthetic variant library based on Omicron BA.1 RBD. **a**, The RBD sequence was split into 11-12 fragments, each being approximately 48 nt in length. For each fragment, a ssODN library with either zero, one or two mutations was designed. **b**, To introduce mutations, NNK codons were tiled across the fragments (1). Each fragment was flanked by BsmBI sites (2). The ssODNs were flanked by primer binding sites for double stranded synthesis through PCR (primers are represented by black arrows and primer binding sites are peach colored) (3). The type II-S restriction enzyme BsmBI gives rise to orthogonal four nt overhangs, which are used by a ligase to assemble individual fragments into full-length RBD sequences (4). **c**, The use of GGA for library construction required the presence of constant regions for ligation between fragments (in black), thereby restricting the library diversity. To overcome this drawback, four staggered sub-libraries were constructed. Due to limitations in sequencing length, it was further necessary to split the RBD into two separate libraries. The extent of seq-library A is indicated in orange and seq-library B in cyan. The primer binding sites for deep sequencing are indicated using orange and cyan arrows. **d**, Targeted sequencing of seq-libraries A and B showed comprehensive mutational coverage for both libraries. The same color scheme as in (c) was used to indicate the extent of both libraries. **e**, To adjust the mutational rate of the library, three different conditions were tested. Different amounts of fragments with zero, one or two mutations were pooled in different ratios which yielded libraries with different mutational distributions.

The current read length of Illumina does not allow coverage of the entire RBD with a single sequencing read (paired-end). Therefore, two separate sequencing libraries (seq-library A and B) were individually constructed. The seq-library A and B possessed mutations in positions 331 - 475 and 386 - 531, respectively (Fig. 1c). The seq-libraries were constructed separately but all subsequent steps were performed in a pooled fashion. Following deep sequencing, complete mutational coverage for each residue was observed in both seq-libraries (Fig. 1d). Interestingly, the mutational frequency is somewhat variable across the seq-libraries, showing a marked decrease in mutations every 16 residues. The low mutational frequencies line up with GGA homologies of sub-library 1. We hypothesize that when pooling the sub-libraries, sub-library 1 was more prominent than the other sub-libraries and therefore less mutations at these sites are observed.

Next, to optimize the number of mutations per RBD sequence, titration of the fragment assembly step was performed. Wild-type (WT) fragments (BA.1 sequence) and fragments with one and two mutations respectively were pooled in different ratios for assembly. Separately, assembly was performed with 60%, 70% and 80% of WT fragments, with the remaining percentage split evenly between fragments with one and two mutations. Deep sequencing of these libraries revealed a clear trend in mutational distribution based on the different ratios, highlighting the tunable nature of our approach. Based on these results, all subsequent work was carried out using the 60% WT library as it has the highest mean number of mutations, therefore providing an appropriate approximation for extensively mutated Omicron sublineages.

### Screening RBD libraries for ACE2 binding and antibody escape

Co-transformation of yeast cells (*S. cerevisiae,* strain EBY100) using the PCR amplified RBD library and linearized plasmid yielded more than 2 x 10^8^ transformants (Methods). Yeast surface display of RBD variants was achieved through C-terminal fusion to Aga2^37^. Next, fluorescence-activated cell sorting (FACS) was used to isolate yeast cells expressing RBD variants that either retained binding or completely lost binding to dimeric soluble human ACE2 (Fig. 2a). Notably, RBD variants with only partial binding to ACE2 were not isolated, as such intermediate populations could not be confidently classified as either binding or non-binding. Removing these variants is essential to obtain cleanly labeled datasets for training supervised machine learning models.

**Figure 2.**
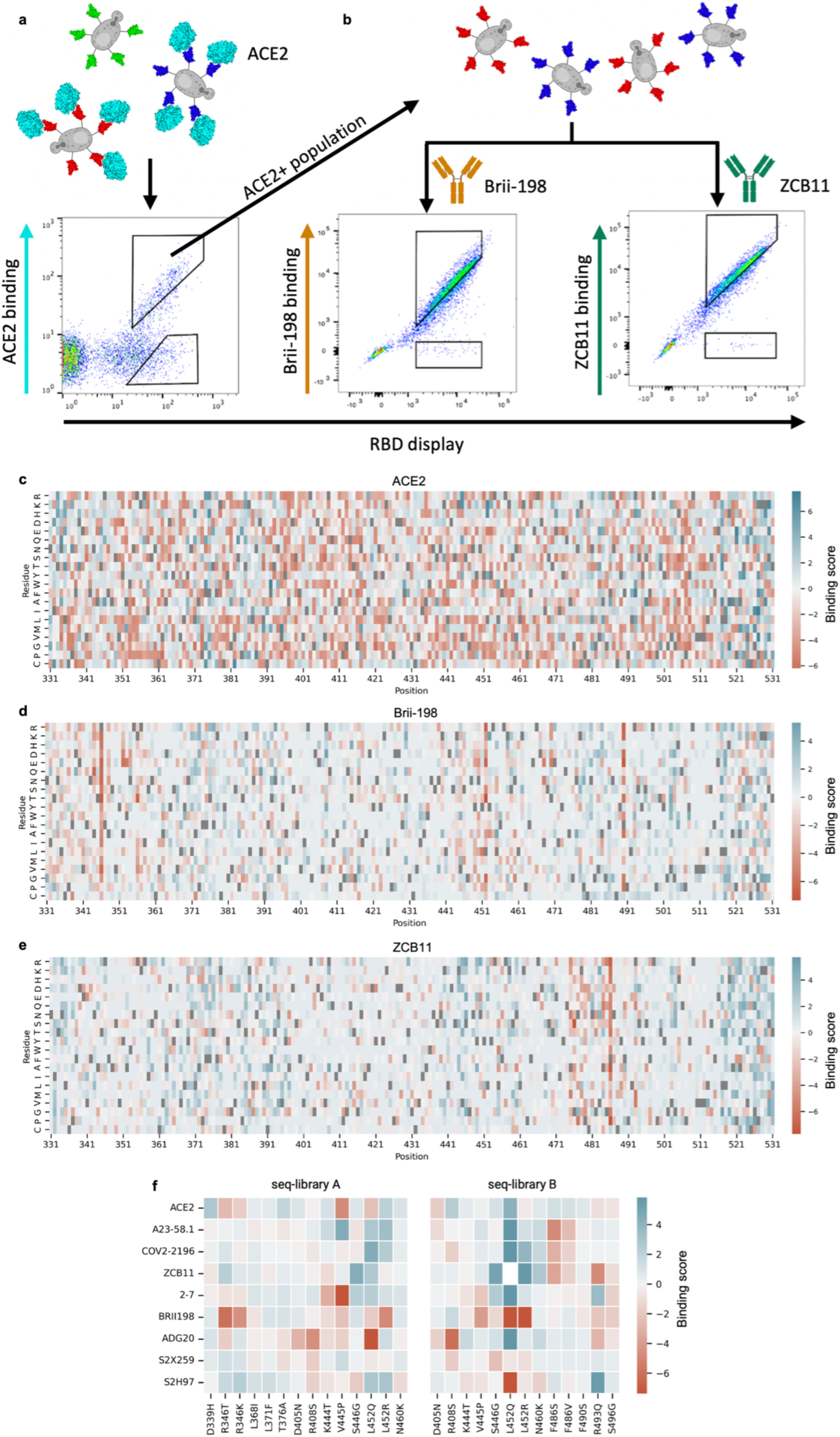
Screening RBD libraries for ACE2 binding and antibody escape by yeast display and deep sequencing. **a**,**b**, Workflow for sorting of yeast display RBD libraries and FACS dot plots for **a,** ACE2 and **b**, antibodies Brii-198 and ZCB11. Gating schemes correspond to binding and non-binding (escape) RBD variant populations. **c**,**d**,**e** Heatmaps depict the binding score of each aa per position of full-length RBD following sorting and deep sequencing of libraries for **c**, ACE2 **d**, Brii-198 **e,** and ZCB11; higher binding score indicates greater frequency in the binding population vs non-binding population. WT BA.1 residues are in gray. **f**, Heatmaps for seq-libraries A and B depict binding scores for ACE2 and antibodies of key mutations seen in major Omicron sublineage variants.

Since binding to ACE2 is a prerequisite for cell entry and subsequent viral replication, only this population is biologically relevant. Thus, only the ACE2-binding population was used in following FACS sorts to isolate RBD variants that either retained binding or completely lost binding (escape) activity to a panel of eight neutralizing antibodies (Fig. 2b, Supplementary Fig. 2 and Supplementary Table 2). The antibodies selected target different epitopes, and are well characterized for their neutralizing activity to BA.1 and its sublineages, which provide a good internal control to assess the accuracy of our method^38–40^. The panel consists of the following antibodies: A23-58.1^41^, COV2-2196^42^, Brii-198^43^, ZCB11^44^, 2-7^45^, S2X259^46^, ADG20^47^, and S2H97^20^.

Following ACE2 and monoclonal antibody sorting, pure populations of RBD variants (binding and non-binding) were subjected to deep sequencing (Supplementary Table 3). Reads covering the RBD sequence were then extracted from the NGS data and heatmaps were constructed depicting binding scores (relative aa frequencies per position in the RBD of binding vs non-binding variants) (Fig. 2c-e and Supplementary Fig. 3). The heatmaps demonstrate nearly complete coverage of mutations across the RBD within all sorted populations. A heterogeneous distribution of mutations is observed for ACE2 binding, with no specific positions or mutations showing dominance (Fig. 2c). This agrees with previous studies that suggest the Q498R and N501Y mutations present in BA.1 exhibit strong epistatic effects that compensate for many mutations that cause loss of binding^48^. In contrast, for certain antibodies, clear mutational patterns could be observed, including escape mutations that correspond with previous DMS studies (Fig. 2d-f and Supplementary Fig. 3). For example, RBD escape variants for Brii-198 are enriched for mutations in positions 346 and 452 (Fig. 2d), which are present in BA.1 and BA.4/BA.5, respectively and correspond to previous work that shows they drive a drastic loss of binding to Brii-198^49^. In contrast, enrichment of these escape mutations are not observed for antibody 2-7 (Supplementary Fig. 3), even though Brii-198 and 2-7 share a similar epitope, suggesting that the binding modality between these two antibodies are different, which is also reflected by their difference in resistance to Omicron variants (e.g., 2-7 shows strong binding to BA.2 and BA.4/BA.5, while Brii-198 does not bind BA.2.12 and BA.4/BA.5)^39,50^. Similarly, the F486V mutation, which has been demonstrated to drastically reduce the neutralization potency of ZCB11 by over 2000-fold^10^, is highly enriched in the RBD escape population (Fig. 2e, f). These mutations are also seen in A23-58.1 and COV2-2196, which bind to a similar epitope (Supplementary Fig. 3). Lastly, for ADG20, we observe a high enrichment of escape mutations in 408(Fig. 2f, Supplementary Fig. 3); this position is also mutated in BA.2 and BA.4/BA.5 variants, which have been shown to have drastically reduced neutralization by ADG20^10^.

While heatmap analysis allows specific mutational patterns to be linked with antibody escape profiles, the high-dimensional nature – and potentially higher order impact – of combinatorial mutations is not reflected in this format. It is apparent that protein epistasis and combinatorial mutations can modify the effect of known escape mutations, either amplifying or reducing antibody binding. For example, individual RBD mutations (G339D, S371F, S373P, S375, K417N, N440K, G446S, S477N, T478K, E484A, Q493R, G496S, Q498R, N501Y, Y505H) in BA.1 and BA.1.1 do not enhance escape to COV2-2196, with each mutation causing an average fold reduction of 2.2, but together cause over 200-fold reduction in neutralization^51^. Conversely, the introduction of the single R493Q mutation in BA.2 substantially rescued the neutralizing activities of Brii-198, REGN10933, COV2-2196 and ZCB11^10^. Thus, while the heatmaps indicate specific mutational contributions to antibody escape, other techniques such as deep learning are required to capture the high-dimensional nature of combinatorial mutations, and generalize to future mutations.

### Deep learning ensemble models accurately predict ACE2 binding and antibody escape

To address the high dimensionality of our dataset and to understand epistatic effects between mutations in the full RBD mutational sequence space, which is far too vast to be comprehensively screened experimentally, we trained deep learning ensemble models. Deep sequencing data from FACS-isolated yeast populations underwent pre-processing and quality filtering prior to being used as training data for machine learning. In the datasets for all antibodies, using the BA.1 RBD sequence as a reference, the mean rate of mutations ranged between ED two (ED_2_) and three ED_3_, with a max ED_8_ (Methods and Supplementary Fig. 5 and 6). Following nucleotide to protein translation, one-hot encoding was performed to convert aa sequences into an input matrix for machine and deep learning models (Fig. 3a). Supervised machine learning models were trained to predict the probability (*P*) that a specific RBD sequence will bind to ACE2 or a given antibody. A higher *P* signifies a stronger correlation with binding, whereas a lower *P* corresponds to non-binding (escape). The machine learning models tested included K-nearest neighbor (KNN), logistic regression (Log Reg), naive Bayes (NB), support vector machines (SVM) and Random Forests (RF). Additionally, as a baseline for deep learning models, a multilayer perceptron (MLP) model was also tested. Finally, we implemented a convolutional neural network (CNN) inspired by ProtCNN^52^, which leverages residual neural network blocks and dilated convolutions to learn global information across the full RBD sequence (Fig. 3a).

**Figure 3.**
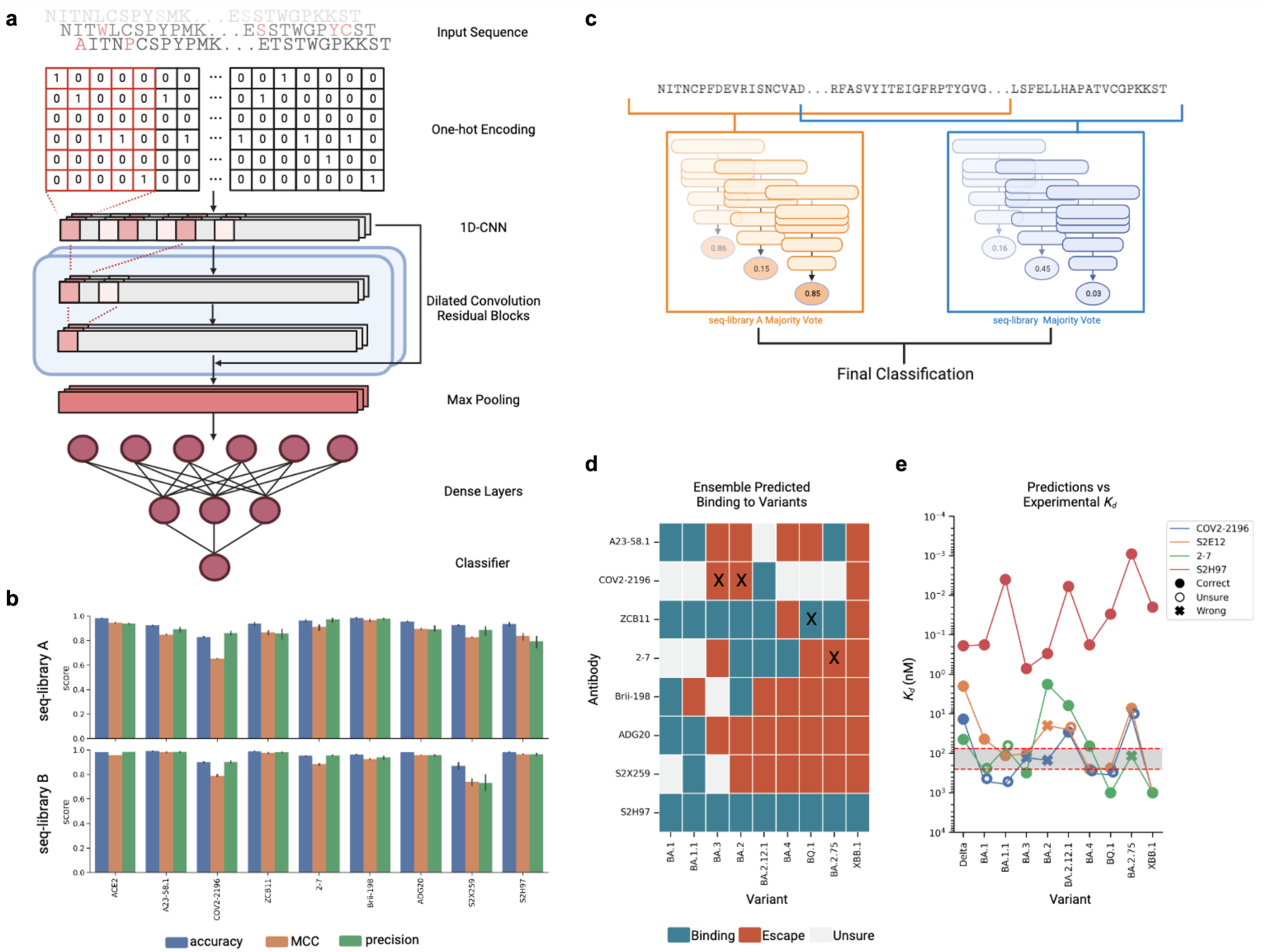
Training and testing of deep learning ensemble models for prediction of ACE2 binding and antibody escape based on full-length RBD sequences. **a**, Deep sequencing data of sorted yeast display libraries are encoded by one-hot encoding and used to train CNN models with several dilated convolutional residual blocks. The models perform a final classification by predicting binding or non-binding to ACE2 or antibodies based on the encoded RBD sequence. **b**, Performance of CNN models trained on all datasets shown by accuracy, Matthews Correlation Coefficient (MCC) and precision. Scores are a result of five rounds of cross-validation with mean performance displayed, and standard deviation indicated by error bars. **c**, Majority voting by an ensemble of models is used to determine the final label for each variant. **d**, Predicted labels of antibodies to well-characterized Omicron variants; colors indicate final labels, and mis-classifications are marked with an “X”. **e**, Comparison of predicted labels to experimental *K_d_* reported in He et al. (2023)^39^ for antibodies 2-7, COV2-2196, S2H97, and S2E12 (as a proxy for A23-58.1), region highlighted in gray indicates model “sensitivity” threshold.

Each model was trained using an 80/10/10 train-validate-test split of data. Inputs were one-hot encoded RBD sequences, with the CNN using a 2D matrix and others using a 1D flattened vector. For initial benchmarking, a collection of different baseline machine learning models were trained on each dataset with hyperparameter optimization through random search, and were evaluated with 5-fold cross validation based on several common metrics (accuracy, F1, MCC, precision and recall). In the baseline machine learning models, class balancing was achieved by random subsampling from the majority class. Unsampled majority class sequences were set aside and merged with the held-out test set for use in model evaluation. Following training, most of the baseline models resulted in relatively high accuracy scores (0.7-0.9) across all datasets, however for smaller datasets (under 20,000 sequences) substantially lower values of F1 (0.2-0.3) and MCC (0.2-0.4) were observed (Supplementary Fig. 7). In contrast, the baseline MLP and CNN deep learning models performed substantially better, including large improvements in F1 and MCC scores (Fig. 3b and Supplementary Fig. 7). While in most cases, the MLP models resulted in relatively high MCC scores (up to > 0.9), CNN models performed substantially better, with MCC scores up to 0.15 higher than MLPs (Supplementary Fig. 7).

Having determined that the CNN models performed superior to the machine learning models and MLP, we next applied an exhaustive hyperparameter search on CNN models to optimize their performance (Supplementary Table 4). Training data was balanced through rejection sampling, while the held-out test set remained imbalanced to accurately evaluate F1 and MCC scores. To prevent data leakage during training, the held-out test set was fixed and multiple models were trained on different training-validation splits of the remaining dataset to make sure each model learned slightly different parameters of the data. When tested on the held-out test set, the final models yielded robust predictive performance up to an ED of eight from the WT BA.1 sequence (Supplementary Fig. 8).

For our final ensemble, we selected three CNN models from each library with the highest MCC scores to generate the predicted labels for each variant through majority voting (Fig. 3c). In short, each model outputs *P* of binding for each input sequence, and labels are assigned based on a threshold. Here, *P* > 0.75 was classified as binding, *P* < 0.25 was classified as non-binding (escape), and those in between were labeled as “uncertain”. The final classification label was taken as the majority label across the three models. An RBD variant was assigned a predicted “escape” label if either the ensemble models of seq-library A or seq-library B predicted escape, and assigned a predicted “binding” label only if both models predicted binding. This leads to a more conservative prediction of antibody binding to variants, and minimizes false-positives. We tested the performance of the ensemble models on published experimental data of antibody binding (or neutralization) to Omicron sublineages^10,38,49,53–56^. In general, the ensemble model predictions performed well, assigning accurate labels to over 80% of the antibody-variant pairs, with only four mis-classifications (Fig. 3d). Three of these mis-classifications were false-negatives, which is likely due to the more conservative approach used for binding classification (Fig. 3d, Supplementary File). Comparing model predictions with published data on antibody affinity values (equilibrium dissociation constant, *K_d_*), revealed that uncertain and mis-classifications were confined to antibodies with intermediate affinities (*K_d_* = 75 - 250 nM), suggesting that there may be a sensitivity limit correlated with lower antibody affinity (Fig. 3e).

### Designing antibody combinations by predicting resistance to synthetic Omicron lineages

After validating the performance of CNN models on test and validation data, we next deployed them to evaluate the resistance of antibodies to viral evolution. While antibody breadth is normally evaluated retroactively based on neutralization or binding to previously observed variants, here we aimed to leverage this machine learning-guided protein engineering approach to prospectively characterize and assess the breadth of antibodies against Omicron variants that may emerge in the future. This was achieved by generating synthetic lineages stemming from BA.1. Since the potential sequence space of combinatorial RBD mutations is exceedingly massive, it was necessary to reduce this to a relevant subspace, therefore mutational probabilities were calculated across the RBD using SARS-CoV-2 genome sequencing data (available on Global Initiative on Sharing Avian Influenza Data, GISAID [www.gisaid.org]) and used to generate synthetic lineages that mimic natural mutational frequencies. Starting with the BA.1 sequence, mutational frequencies from 2021 and 2022 were utilized to generate ten sets of 250,000 synthetic RBD sequences through six rounds of *in silico* evolution, where the 100 variants with the highest predicted score for ACE2 binding (averaged across the ensemble CNN models) in each round were used as seed sequences for the next round of mutations. Next, the ensemble deep learning models were used to predict antibody binding or escape (or uncertain classification) for the synthetic variants. This provides an estimation of each individual antibody’s binding breadth in the generated sequence space and thus correlates with resistance to prospective Omicron lineages (Fig. 4a,b, Supplementary Fig. 9).

**Figure 4.**
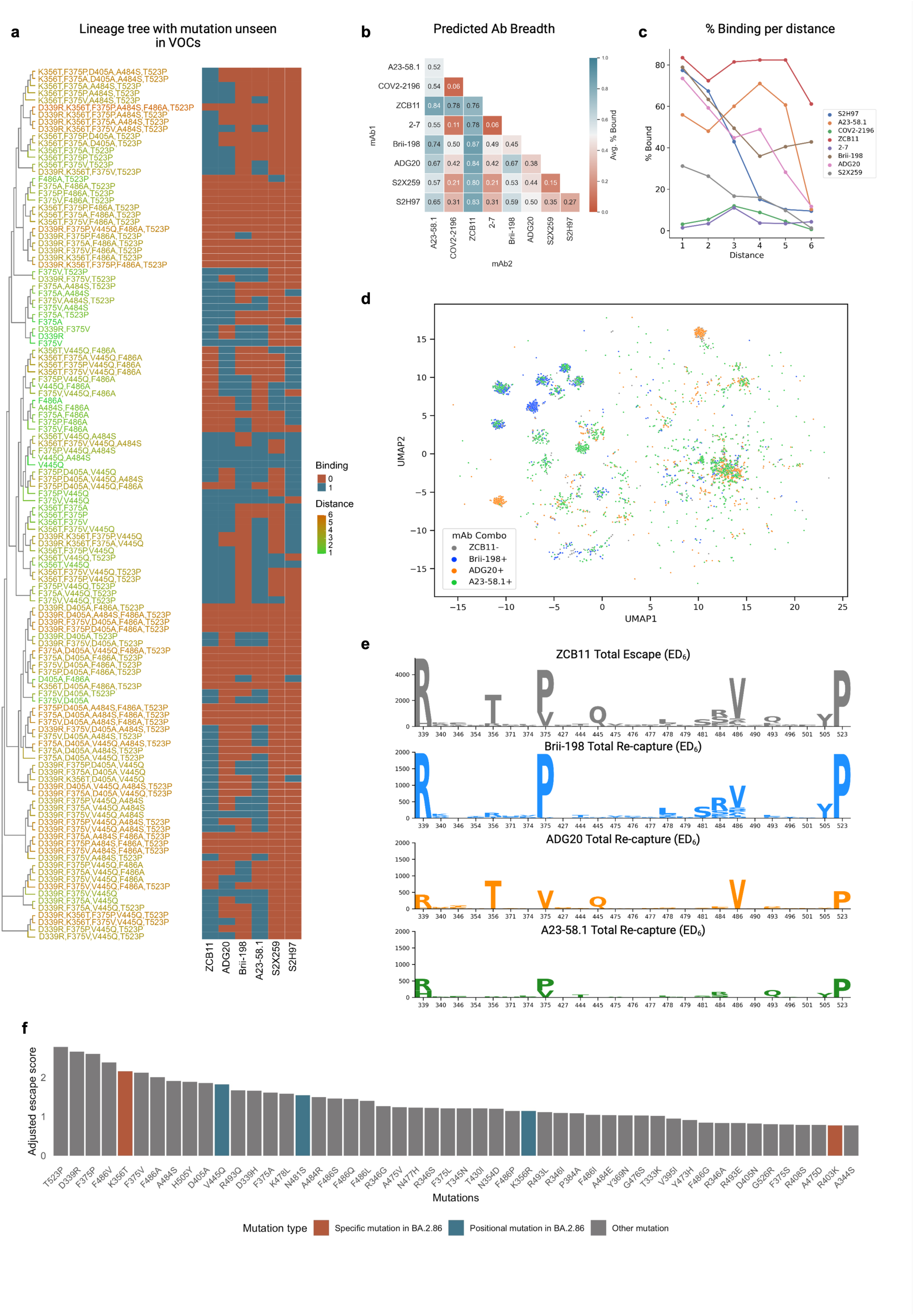
Evaluating antibody breadth on synthetic Omicron lineages. **a**, Example of a synthetic lineage tree of sequences generated containing mutations unseen in major Omicron variants, with heatmap indicating the deep learning predictions of binding or escape for individual antibodies. **b**, Total mean predicted breadth of individual antibodies and combinations on synthetic lineages generated from 2022 mutational probabilities. **c**, The fraction (%) of sequences bound by individual antibodies at different ED from BA.1. **d**, UMAP displays a subsample of ZCB11 escape variants in protein sequence space with antibody-specific binding clusters highlighted. **e**, Sequence logos show the top 25 positions with greatest Kullback-Leibler (KL)-divergence in ZCB11 escape variants at ED_6_, and sequences re-captured by Brii-198, ADG20 and A23-58.1. **f**, The top 50 predicted mutations ranked by their escape scores (see Methods) from the generated synthetic lineages, with new mutations seen in the BA.2.86 variant highlighted.

Since several of the clinically used antibody therapies for COVID-19 consisted of a cocktail of two antibodies (e.g., LY-CoV555+LY-CoV16, REGN10933+REGN10987, COV2-2130+COV2-2196), we also determined antibody breadth across all two-way combinations. For the 2022-based synthetic lineages, ZCB11 showed the greatest predicted breadth, followed by A23-58.1, Brii-198 and ADG20 (Fig. 4b). The ensemble models predict very low breadth for 2-7 and COV2-2196, despite both maintaining binding to BA.2 and beyond^39^. This is likely due to the high uncertainty of these models. The predicted coverage of ZCB11 corresponds well with experimental measurements that show it maintains high affinities and neutralization to several Omicron variants (BA.2, BA.4/5)^10^. Similarly, Brii-198 and A23-58.1 have been shown to bind BA.2, BA.2.12 and BA.2.75 variants^40^, aligning with the predictions of their relatively high breadth. Examining breadth profiles of each antibody as a function of ED revealed differing profiles, such as ZCB11 and Brii-198 maintaining high breadth at larger ED (>ED_4_), while A23-58.1 and ADG20 have substantially lower breadth at large ED (Fig. 4c). The predicted breadth of several antibodies were substantially different for synthetic lineages generated using 2021 mutational probabilities. For example, the breadth of ADG20 is substantially higher as it is predicted to bind over 50% of variants, while the breadth of Brii-198 and A23-58.1 is reduced by 9% and 15%, respectively (Supplementary Fig. 9 and 10). This suggests that correctly anticipating antigenic drift and changes in mutational frequencies play an important role in determining breadth predictions.

It is worth noting that calculating the breadth of antibody combinations is not simply additive. For example, while Brii-198 ranks lower than A23-58.1 in total breadth, Brii-198 provides more complementary coverage to ZCB11 (Brii-198 binds to more variants that escape ZCB11), resulting in an overall increase in variant coverage in a simulated cocktail. Examining the distribution of escape variants for ZCB11 at ED_6_, where it sees its most significant breadth reduction— the three other highly ranked antibodies (A23-58.1, Brii-198 and ADG20) re-establish coverage over unique clusters in the sequence space (Fig. 4d). However, only ADG20 and Brii-198 cover and mitigate variants that include the key F468V mutation (e.g., BA.4/5). Furthermore, Brii-198 covers the most diverse clusters that contain additional critical mutations at the F468 position, in addition to the surrounding residues in this epitope (Fig. 4e and Supplementary Fig. 11). Thus, while any of the three antibodies would be complementary to ZCB11 by nature of targeting a different epitope^10^, our breadth analysis aids in identifying the most complementary antibody by variant coverage.

To quantify the impact of how individual mutations can drive antibody escape, an escape score (*S*_*m*^) was computed for each mutation (*m*) within the synthetic lineages. This metric is a normalized product of the number of antibodies escaped by a given mutation and the mutation’s frequency within the lineage (see Methods). When examining individual RBD mutations across the synthetic lineages (Fig. 4f), it was revealed that T523P has the highest escape score. Comparatively, DMS results showed that mutations at position 523 have a slightly negative influence on RBD protein expression level ^19^, which may explain its low occurrence in natural variants, having only been observed in 70 sequences in the GISAID database. Furthermore, the combination of D339R, F486A and T523P mutations in the simulated BA.1 lineages caused the most antibody escape among mutations not previously observed in major variants (Fig. 4f). Out of these, the positions 339 and 486 are mutated in BA.2.75 and XBB and their sublineages. The top 50 mutations with the highest escape scores include K356T and R403K, which are present in the recently reported and highly mutated BA.2.86 variant and had not been previously reported in any other major variant (Fig. 4f). Additionally, positions V445 and N481 were also mutated in BA.2.86. Taken together, this suggests that DML-derived escape scores may reveal mutations or positions that emerge in future variants.

## DISCUSSION

The emergence of SARS-CoV-2 lineages with a high number of mutations has resulted in substantial viral immune evasion, including ineffective neutralization by previously developed therapeutic antibodies^5^. This rapid pace of viral evolution has underscored the need for novel approaches to adequately profile antibody candidates and predict their robustness to emerging variants early on during drug development. To this end, we leverage DML, a machine learning-guided protein engineering method to prospectively evaluate clinically relevant antibodies for their breadth against potential future Omicron variants across a large mutational sequence space.

We first demonstrate the feasibility of assembling full-length RBD mutagenesis libraries with high fidelity using a large number of relatively short ssODNs in a one-pot reaction and obtaining library sizes in excess of 10^8^. This is despite the fact that previous studies have reported a decrease in GGA when increasing the number of DNA fragments^39^. Screening of these libraries for ACE2 binding and antibody escape yielded high-dimensional data sets with combinatorial mutations spanning the entire RBD sequence, which is not obtainable through frequently employed approaches such as DMS. In addition, the RBD library design can be updated to accommodate mutations present in emerging variants, and the average number of mutations can be titrated to generate data suitable for the training of machine learning models. This library design and screening approach could also be exploited to profile viral surface proteins from other rapidly evolving viruses such as influenza or HIV, two viruses which undergo substantial antigenic drift that drives their immune escape^57–59^.

So far, the breadth of SARS-CoV-2 therapeutics has been assessed through the use of past variants and observed mutations^20,60–62^. Measuring breadth in this way does not adequately predict long-term resistance against future variants. The deployment of ensemble deep learning models to make predictions on synthetic mutational trajectories of the RBD enabled an effective quantitative method to evaluate the breadth of each antibody based on its coverage of RBD mutational sequence space. DML predictions confirm that ZCB11 has exceptionally broad breadth to major Omicron lineages that emerged in 2022, while many other antibodies fail against Omicron variants^39^. Furthermore, our results suggest that the standard structure-based approach of selecting antibodies targeting different epitopes in a cocktail does not sufficiently determine which combinations offer the most cumulative breadth. High breadth cocktails would ensure that even if a variant escapes one antibody in the cocktail, it has a high chance to be re-captured by the other antibody - thus potentially maintaining the clinical effectiveness of the therapy. For example, this occured with the combination antibody therapy from Eli Lilly (LY-CoV555+LY-CoV16), which continued to be used clinically when only a single antibody in the combination was effective after the emergence of Beta, Gamma and Delta variants^22,63^. Interestingly, a comprehensive search through a SARS-CoV-2 antibody database (Cov-AbDab, accessed April 2023)^64^ reveals that a number of neutralizing antibodies discovered early in the pandemic from patients infected with the ancestral Wu-Hu-1 are still able to neutralize Omicron variants such as BA.5, BQ.1 and XBB.1. DML could therefore be a powerful tool to identify such variant-resistant antibodies for therapeutic development.

Analysis of DML breadth predictions also highlights specific and positional mutations that are associated with greater immune escape, with four such mutations being observed in the recently discovered and highly mutated BA.2.86 variant. In contrast, other recently published deep learning methods, which rely on models trained using a combination of DMS and protein structure data, were able to only correctly forecast one new mutation each that appeared in the XBB.1.5 and BQ.1 variants, respectively^30,31,65^. While this demonstrates the value of using protein structural information to better infer higher-order effects between mutations, these models are still limited by the use of low-distance (most often single-mutation) DMS data. Thus, it would be worthwhile to explore whether the use of combinatorial DML data can further improve the accuracy and forecasting performance of models trained using a multi-task objective, similar to those mentioned above.

The accuracy of antibody breadth predictions is dependent on having an accurate forecast of future mutations in the RBD. The use of deep learning models that predict ACE2 binding allowed us to capture evolutionary pressures correlated with host receptor binding, which is a mandatory feature of any emerging SARS-CoV-2 variant^66^. However, a myriad of other factors impact antigenic drift and variant emergence, such as transmissibility, host cell infectivity, crossover, reproductive rate, etc.^67^, thus generating training data related to these factors, for example through the use of an advanced pseudovirus mutational library screening system^68^, may further support the generation of deep learning models that can predict future mutations and variants with higher accuracy.

## METHODS

### Construction of a high distance Omicron RBD library for yeast surface display

Synthetic ssODNs (oPools from IDT) were designed with either one or all possible combinations of two degenerate NNK codons for each fragment (Supplementary Table 1). For each fragment, 136 ssODNs were designed (16 single NNK codons and 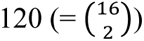 double NNK codon combinations). Each fragment was flanked by BsmBI recognition sites and ∼20 nt for second strand synthesis through PCR. For high fidelity library assembly, the overhangs were optimized using the NEB ligase fidelity viewer (https://ligasefidelity.neb.com/viewset/run.cgi). Using the NEBridge^®^ Golden Gate Assembly Kit (NEB, E1602), individual fragments were assembled to full-length RBD gene segments. A custom entry vector based on pYTK001 (addgene, Kit #1000000061) was designed. Double stranded fragments were mixed with 75 ng entry vector in a 2:1 molar ratio. As suggested by the manufacturer’s instructions, 2 μL NEB Golden Gate Enzyme Mix was used. For the assembly, the following protocol was used: (42°C, 5 min → 16°C, 5 min) x 30 → 60°C, 5 min. The assembled libraries were transformed into *E. coli* DH5α ElecroMAX (Thermo Fisher Scientific, 11319019), resulting in ∼4 x 10^8^ transformants. According to the manufacturer’s instructions (Zymo, D4201), the RBD library plasmid was extracted from *E. coli*.

The RBD library was PCR amplified and the yeast display vector (pYD1) was linearized using the restriction enzyme BamHI (Thermo Fisher Scientific, FD0054). Both insert and backbone were column purified according to the manufacturer’s instructions (D4033) and drop dialyzed for 2 h using nuclease-free water (Millipore VSWP02500). The RBD library insert and linearized pYD1 backbone were co-transformed into yeast (*S. cerevisiae,* strain EBY100) using a previously described protocol^69^. Briefly, EBY100 (ATCC, MYA-4941) was grown overnight in YPD [20 g/L glucose (Sigma-Aldrich, G8270), 20 g/L vegetable peptone (Sigma-Aldrich, 19942), and 10 g/L yeast extract (Sigma-Aldrich, Y1625) in deionized water]. On the day of the library preparation, yeast cells from the overnight culture were inoculated in 300 mL YPD at an OD_600_ of 0.3. The cells were grown to an OD_600_ of 1.6 before washing the cells twice with 300 mL ice cold 1 M Sorbitol solution (Sigma-Aldrich, S1876). In a subsequent step, the cells were conditioned using a solution containing 100 mM lithium acetate (Sigma-Aldrich, L6883) and 10 mM DTT (Roche, 10197777001) for 30 min at 30 °C. This was followed by a third wash using 300 mL ice cold 1 M Sorbitol solution. Using 50 μg insert and 10 μg pYD1 backbone, electrocompetent EBY100 were transformed using 2 mm electroporation cuvettes (Sigma-Aldrich, Z706086). The cells were recovered for 1 h in in recovery medium (YPD:1 M Sorbitol solution mixed in a 1:1 ratio) before passageing the cells into selective SD-CAA medium [20 g/L glucose (Sigma-Aldrich, G8270), 8.56 g/L NaH_2_PO_4_·H_2_O (Roth, K300.1), 6.77 g/L Na_2_HPO_4_·2H_2_O (Sigma-Aldrich, 1.06580), 6.7 g/L yeast nitrogen base without amino acids (Sigma-Aldrich, Y0626) and 5 g/L casamino acids (Gibco, 223120) in deionized water]. The cells were grown for 2 days at 30 °C. To estimate the transformation efficiency, dilution plating was performed. Approximately 2 x 10^8^ transformants were obtained.

### Screening RBD libraries for ACE2-binding or non-binding

Yeast cells containing the RBD library plasmid were grown in SD-CAA for 18 - 24 h at 30°C. Surface display of Omicron RBD was induced by passageing the cells into SG-CAA medium [20 g/L galactose (Sigma-Aldrich, G0625), 8.56 g/L NaH_2_PO_4_·H_2_O, 6.77 g/L Na_2_HPO_4_·2H_2_O, 6.7 g/L yeast nitrogen base without amino acids and 5 g/L casamino acids in deionized water]. The cells were incubated at 23°C for 48 hours, as previously described^37^. Approximately 10^9^ cells were spun down by centrifugation at 3500 x g for 3 min and washed once with 5 mL cold wash buffer [DPBS (PAN Biotech, P04-53500)+0.5% BSA (Sigma-Aldrich, A2153)+2 mM EDTA (Biosolve, 051423)+0.1% Tween20 (Sigma Aldrich, P1379)]. Next, cells were labeled with 50 nM of biotinylated human ACE2 protein (Acro Biosystems, AC2-H82E6) for 30 minutes at 4°C at 700 RPM on a shaker (Eppendorf, ThermoMixer C). The cells were subsequently washed. In a secondary staining step, cells were labeled with Streptavidin-Phycoerythrin (PE) (Biolegend 405203) (1:80 diluted) and anti-FLAG Tag Allophycocyanin (APC) (Biolegend 637308) (1:200 dilution) at 4°C for 30 min at 700 RPM. Afterwards, cells were centrifuged at 3500 x g for 3 min. The supernatant was discarded and the tube was protected from light and stored on ice until sorting. Binding (PE+/APC+) and non-binding (PE-/APC+) populations of yeast cells were collected by FACS (BD FACSAria Fusion or BD Influx) (Fig. 2a,b and Supplementary Fig. 2). Collected cells were pelleted at 3500 x g for 3 min to remove the FACS buffer. The cells were resuspended using SD-CAA and grown for two days at 30°C. The sorting process was repeated until the desired populations were pure.

### Screening RBD libraries for antibody binding or escape

The ACE2-binding population of yeast cells expressing the RBD library was grown and induced as described above. Approximately 10^8^ cells were pelleted by centrifugation at 3500 x g for 3 min at 4°C and washed once with 1 mL wash buffer. The washed cells were incubated with antibodies (concentrations listed in Supplementary Table 2). Suitable concentrations approximately corresponding to the EC_90_ were experimentally determined beforehand (Supplementary Fig. 12). Cells were incubated for 30 min at 4 °C and 700 RPM. After an additional washing step, a secondary stain was performed using 5 ng/ml anti-human IgG-AlexaFluor647 (AF647) (Jackson Immunoresearch, 109-605-098) (1:200 dilution). The cells were incubated for 30 minutes at 4 °C and 700 RPM. Subsequently, cells were washed and stained in a tertiary staining step using 1 ng/ml anti-FLAG-PE (1:200 dilution) for 30 min at 4 °C and 700 RPM. Cells were pelleted by centrifugation at 3500 x g for 3 min at 4 °C. The supernatant was discarded and the tube was protected from light and stored on ice until sorting. Cells expressing RBD that maintained antibody-binding (AF647+/PE+) or showed a complete loss of antibody binding (AF647-/PE+) were isolated using FACS (BD Aria Fusion or Influx BD). Collected cells were pelleted by centrifugation at 3500 x g for 3 min at room temperature. The FACS buffer was discarded and the cells were resuspended using SD-CAA. The cells were cultured for 48 h at 30 °C. The sorting process was repeated once for the binding population and twice for the non-binding population. This procedure yielded pure binding and non-binding (escape) populations.

### Deep sequencing of RBD libraries

The pYD1 plasmid encoding the RBD library was extracted from yeast cells per manufacturer’s instructions (Zymo, D2004). The mutagenized part of the RBD was PCR amplified using custom designed primers for seq-library A and seq-library B (Supplementary Table 5). In a second PCR amplification step, sample specific barcodes (Illumina Nextera) were introduced, which allowed pooling of individual populations for sequencing. The populations were sequenced using the Illumina MiSeq v 3 kit which allows for 2 × 300 paired-end sequencing.

### Preprocessing of deep sequencing data

Sequencing reads were paired, quality trimmed and merged using the BBTools suite^70^ with a quality threshold of qphred R>25. RBD nt sequences were then extracted using custom R scripts, followed by translation to aa sequences. Read counts per sequence were calculated and singletons (read count = 1) were discarded. Sequencing datasets used for training machine and deep learning models were created by combining the binding and non-binding datasets. Sequences present in both populations were removed.

Binding scores for heatmaps shown in Fig. 2c-e were created by calculating aa counts per position in the RBD from both binding and non-binding sequences. WT (BA.1) aa residues were then removed, relative frequencies were calculated with a pseudocount of 1 added, and final binding scores were calculated as binding frequencies divided by non-binding frequencies. The results were then log-transformed before plotting in the heatmap for visualization.

### Training and testing machine and deep learning models

All machine learning code and models were built in Python (3.10.4)^71^. For data processing and visualization, numpy (1.23.3), pandas (1.4.4), matplotlib (3.5.3) and seaborn (0.12.0) packages were used. Baseline benchmarking models were built using Scikit-Learn (1.0.2), while Keras (2.9.0) and Tensorflow (2.9.1) were used to build the MLP and CNN models.

Each model was trained using 80/10/10 train-val-test data random splits. RBD library protein sequences (from seq-library A or B deep sequencing data) were one-hot encoded prior to being used as inputs into the models. For the CNN, the 2D one-hot encoded matrix was used as the input, while for others, the matrix was flattened into a one-dimensional vector. All reported model performances were evaluated using 5-fold cross-validation, and evaluated based on the metrics for accuracy, f1, MCC, precision, and recall.

When training baseline machine learning models, class balancing was performed through random downsampling from the majority class so that it was equal to the counts from the minority class; this was performed at each ED. RBD sequences that were not sampled from the majority class were then reserved separately as additional ‘‘unseen sequences’’. These were then combined with the held-out test set during model evaluation to ensure that the models could perform well with an imbalanced test set. Hyperparameter optimization was performed during model training using up to 30 rounds of RandomSearchCV (from Scikit-Learn), and the best model performances were kept for comparison to deep learning models.

To train the deep learning models, exhaustive hyperparameter search was performed on the CNN models to optimize performance through the hyperparameters listed in (Supplementary Table 4). The training dataset was balanced at different ratios (see Minority Ratio row, Supplementary Table 3) while validation and test sets remained unbalanced to appropriately evaluate MCC, precision and recall scores on imbalanced data. Dataset balancing was performed through rejection sampling using a custom dataset sampler created in Tensorflow. To prevent data leakage during training of the models for ensembles, the held-out test set was fixed, while multiple models were trained on random splits of the training and validation sets to make sure each model learned slightly different parameters of the dataset, while being evaluated on the same held-out test sequences.

### Predictions made with ensemble deep learning models

Natural and *in silico* generated synthetic RBD variant sequences were assigned “binding”, “escape” and “uncertain” labels for ACE2 and antibodies using an ensemble of trained models. For a given RBD sequence, each model assigns a binding label if output *P* > 0.75, escape if output *P* < 0.25, or uncertain otherwise. For each of the two libraries (seq-library A and B), the three models with the highest MCC scores were used to independently assign labels to each sequence, followed by majority voting, where the most common label was taken as the label for each variant. The labels from models trained with seq-library A or seq-library B were used to determine the final label for each variant: “binding” if both libraries agree on a “binding” label, “escape” if either library predicts “escape”, and “unsure” otherwise. For experimentally measured variants, antibodies-variant pairs were labeled as “escape” if their measured *K_d_* was > 100nM or IC50 was > 1ug/mL.

### Calculating mutational probabilities of the RBD based on SARS-CoV-2 genome data

To generate the mutational probability matrices used for synthetic lineages, SARS-CoV-2 spike protein sequences were obtained from the GISAID database (most recent access of June 2023) The regions corresponding to the RBD were extracted, along with the date when each sequence was deposited into the database. Sequences were separated by the year they were added (e.g., 2021 or 2022). From these sequences, mutations were counted at each position per position, and per aa. Mutational frequencies at each position were calculated using these counts. Finally, a log softmax function was applied to obtain mutational probabilities for each position. For each position, only residues that were observed in GISAID sequences were counted, while all unseen residues were not included in the softmax transform, preventing them from being generated in synthetic lineages.

### In silico generation of synthetic Omicron lineages

Using BA.1 as the initial seed variant, *in silico* sequences were generated in a stepwise fashion over six rounds of mutations. In the first round, single mutations were randomly generated across the RBD. Positions and aa for each mutational round were selected using probabilities from the 2021 or 2022 substitution matrices; as a control, sequences were also generated using no substitution matrix (where all mutations were sampled from a uniform probabilities distribution). Then binding probability scores were assigned to variants in each generation by taking the average of all *P* predicted by each of the ACE2 models in the ensemble. The top 100 variants ranked by ACE2-binding *P* were used as seed sequences for the next round of mutations. For each round, new variants were only accepted if they contained mutations not previously seen in other generated variants, or else the process was repeated again and new mutations selected until the maximum number of variants were reached (250,000).

### Calculating escape scores

n An escape score (*S_m_*) was calculated that aims to quantify the impact of a given mutation on driving escape from the antibodies tested herein and was calculated by:

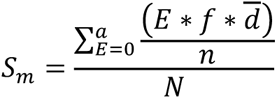

*S_m_* is the escape score of a mutation *m*, *E* is the number of antibodies that are predicted to escape from *m*, and within the group of sequences with the same number of *E*, *f* is the frequency that *m* appeared in the sequence group, <u>*d*</u> is the mean of sequence ED from BA.1, *n* is the number of sequences, *N* is how many times one mutation appeared in different groups of *E, a* is the total number of groups, according to how many antibodies were tested (here, *a* = 6). For better visualization, the adjusted escape score was used (Fig 4a, f) and is calculated by the following equation:

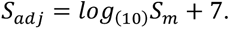

### Additional statistical analysis and plots

Statistical analysis was performed using Python (3.10.4) with the Scipy package (1.9.3). Dimensionality reduction was performed using UMAP-learn (0.5.3). Graphics were generated using matplotlib (3.5.3), seaborn (0.12.0), and ggtree (3.8.0). Sequence logo plots were created using Seq2Logo (5.29.8)^72^ or the dsmlogo package from the Bloom Lab (https://github.com/jbloomlab/dmslogo).

The KL-divergence was calculated by adapting a recently described method^30^. In short, a probability-weighted KL logo plot was used to visualize differences between a subset of sequences to the full background dataset. Let M1 = (f_1_, f_2_, f_3_…, f_n_) represent the position frequency matrix (PFM) of the background sequence set, where the length of the initial sequence is n = 201 and each frequency f_i_ = (*a_1_, a_2_, a_3_,…a_20_*)^T^, represents the frequency of each aa per position i. At the same time, M2 = (f_1_′, f_2_′, f_3_′,…f_n_′) represents the PFM of the subset of sequences, each f_i_′ = (*a_1_′, a_2_′, a_3_′,…a_20_′*)^T^. The KL divergence at each position is computed as:

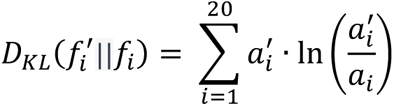

The KL divergence is used to set the total height at each position in the logo plots (Fig. 4e). The height and direction of each aa letter are calculated through probability-weighted normalization as part of the Seq2Logo package using:

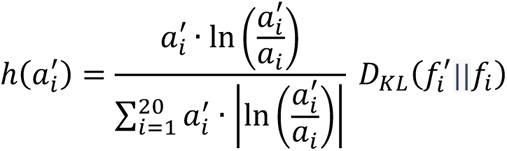

## Data availability

The main data supporting the results in this study are available within the paper and its Supplementary Information. The raw and analysed datasets generated during the study will be made available at: https://github.com/LSSI-ETH/Omicron_DML.

## Code availability

The code and models used to perform the work in this study will be available at the following: https://github.com/LSSI-ETH/Omicron_DML.

## Supporting information

Supp_Table_antibody_data

## Acknowledgments

We thank the ETH Zurich D-BSSE Single Cell Unit and the ETH Zurich D-BSSE Genomics Facility for support. This work was supported by the Botnar Research Centre for Child Health (FTC COVID-19, to S.T.R.)

## Author contributions

L.F., B.G., J.H., J.M.T and S.T.R. developed the methodology. L.F., J.M.T. designed and generated mutagenesis libraries, L.F., performed screening experiments, B.G. and J.H., analyzed the sequencing data and performed deep-learning analyses. L.F., B.G., J.H. and S.T.R. wrote the manuscript, with input from all other authors.

## SUPPLEMENTARY FIGURES

**Supplementary Figure 1.**
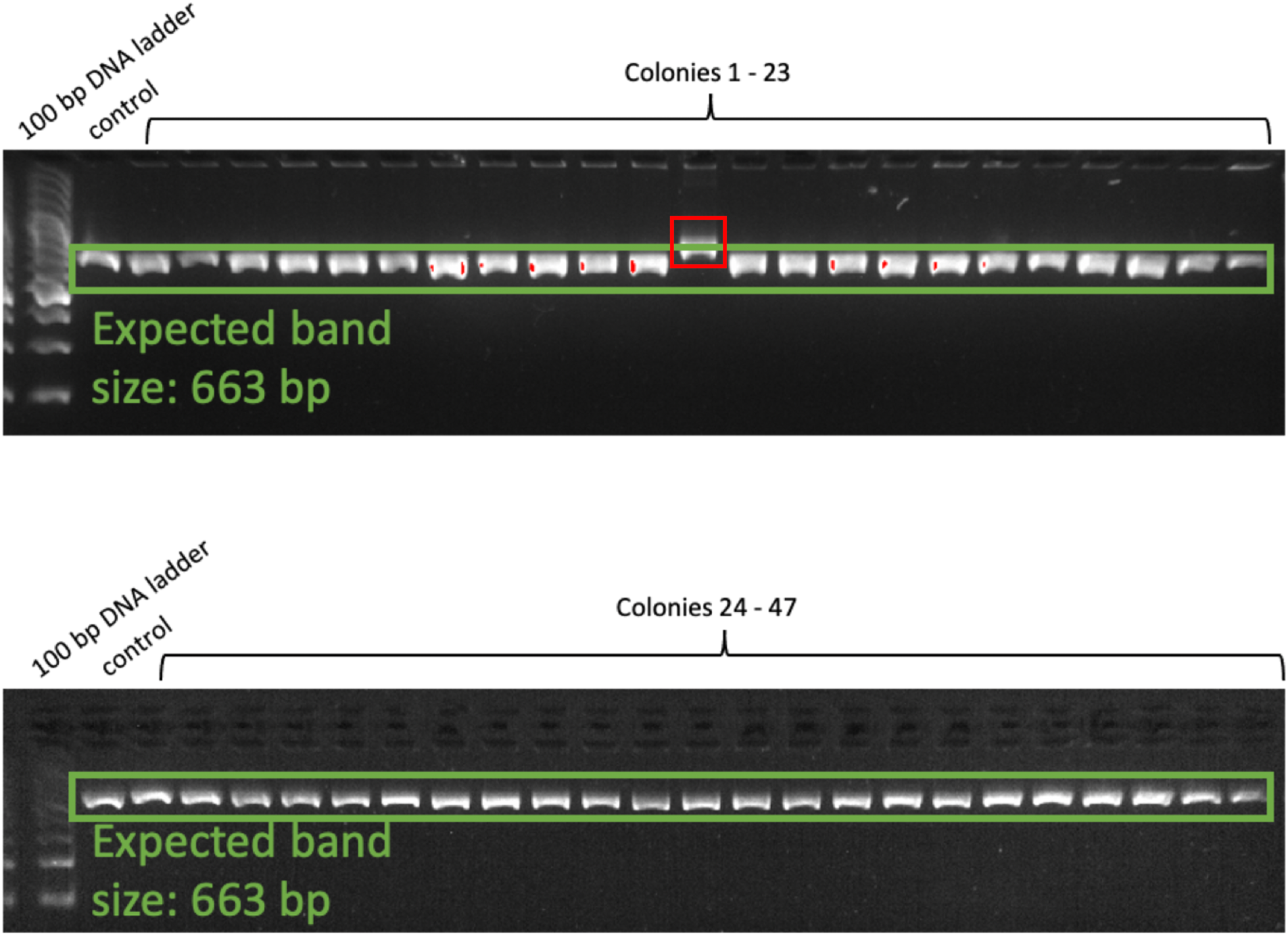
After assembling the RBD sequence from short fragments and transformation into *E. coli*, single colonies were picked and colony PCRs (cPCR) were performed. For the amplification, primers binding directly upstream and downstream of the RBD were used. As a control, WT BA.1 plasmid was used. When running the cPCR products on a 2% agarose gel, 46 out of 47 reactions showed the right band size of 663 base pairs (bp) (wrongly assembled variant highlighted in red), roughly corresponding to 98% correctly assembled full length RBD sequences.

**Supplementary Figure 2.**
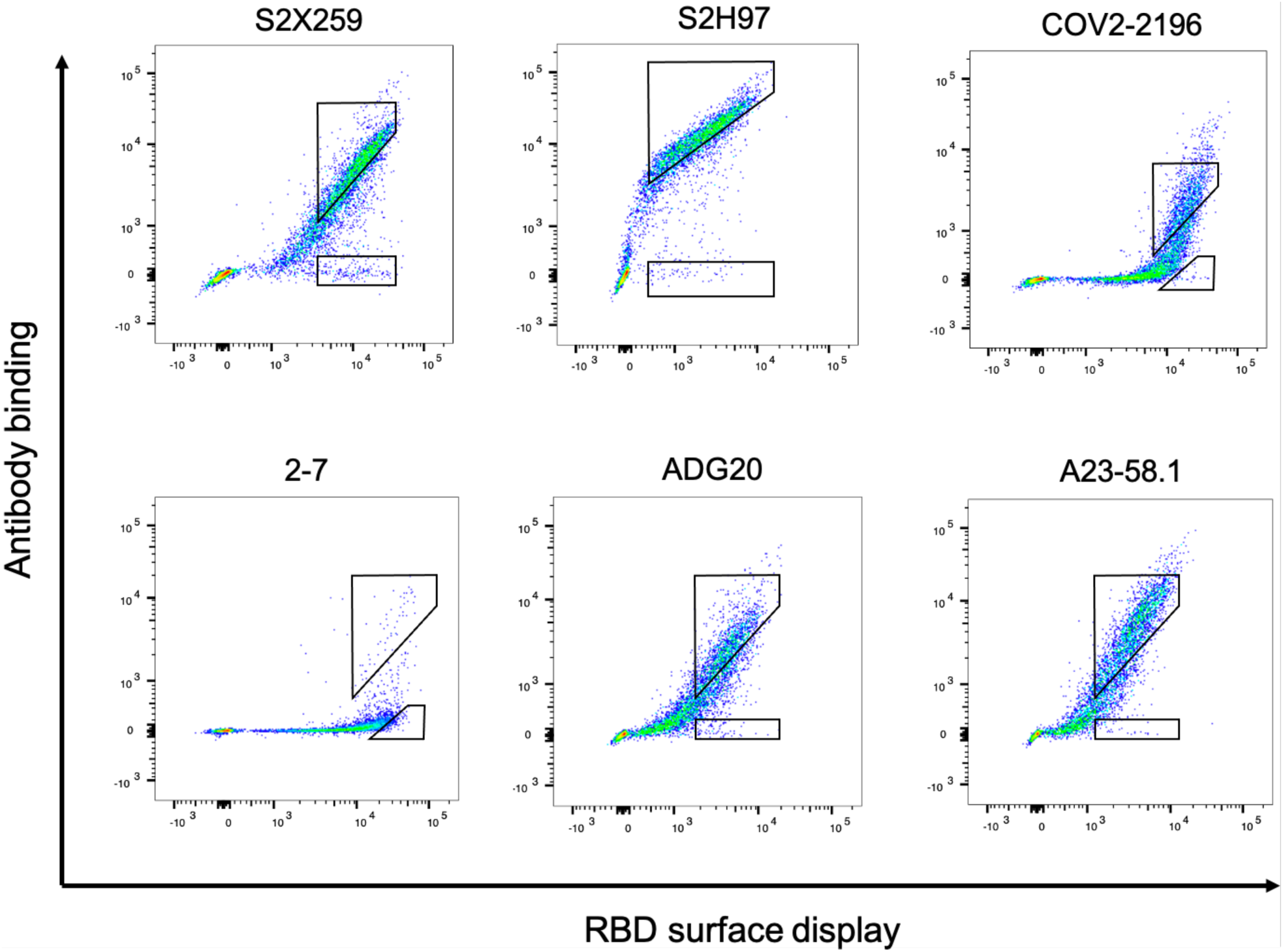
Representative FACS dot plots of yeast RBD libraries during antibody screening; sorting gates for binding and non-binding (escape) populations are shown.

**Supplementary Figure 3.**
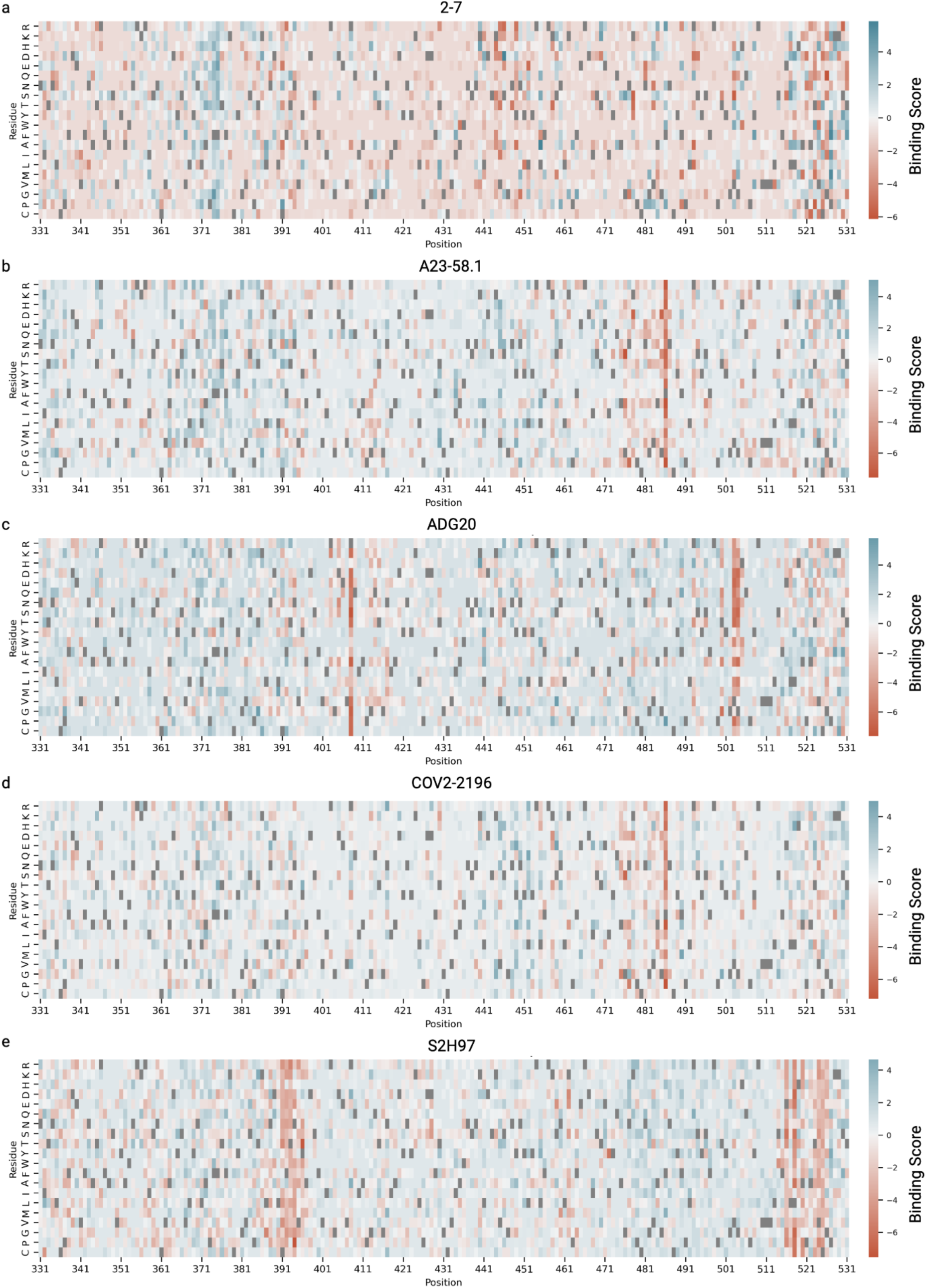
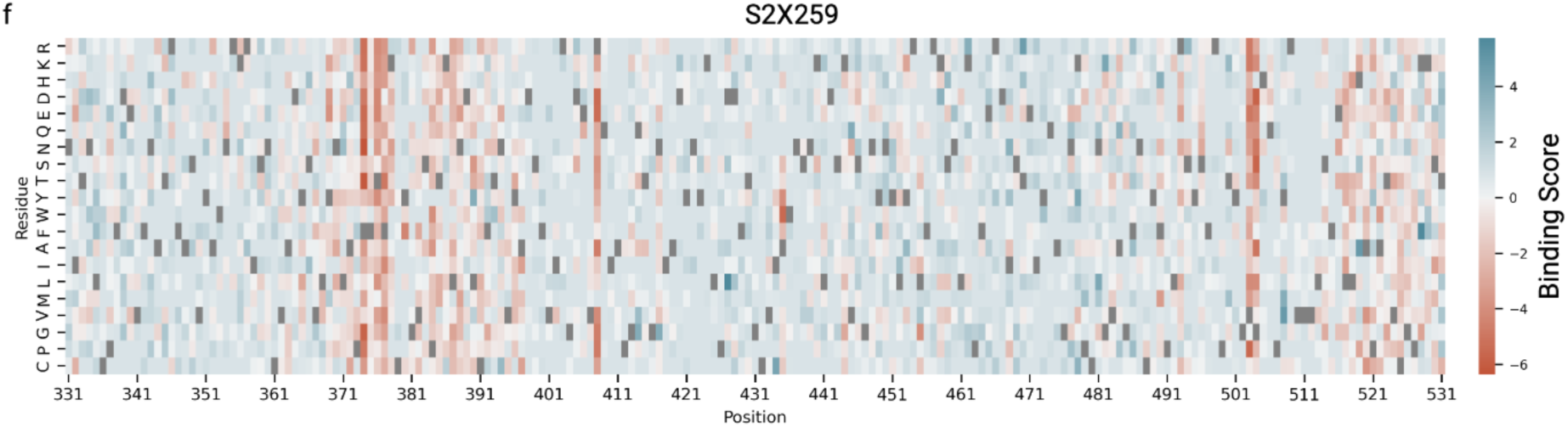
Heatmaps showing binding scores per position across the RBD for libraries sorted against each target (ACE2 or antibodies, respectively). Blue regions indicate mutations seen in greater frequency in the binding variant pool, while red regions indicate mutations with greater frequency in escape variants. WT (BA.1) residues are depicted by grey boxes.

**Supplementary Figure 4.**
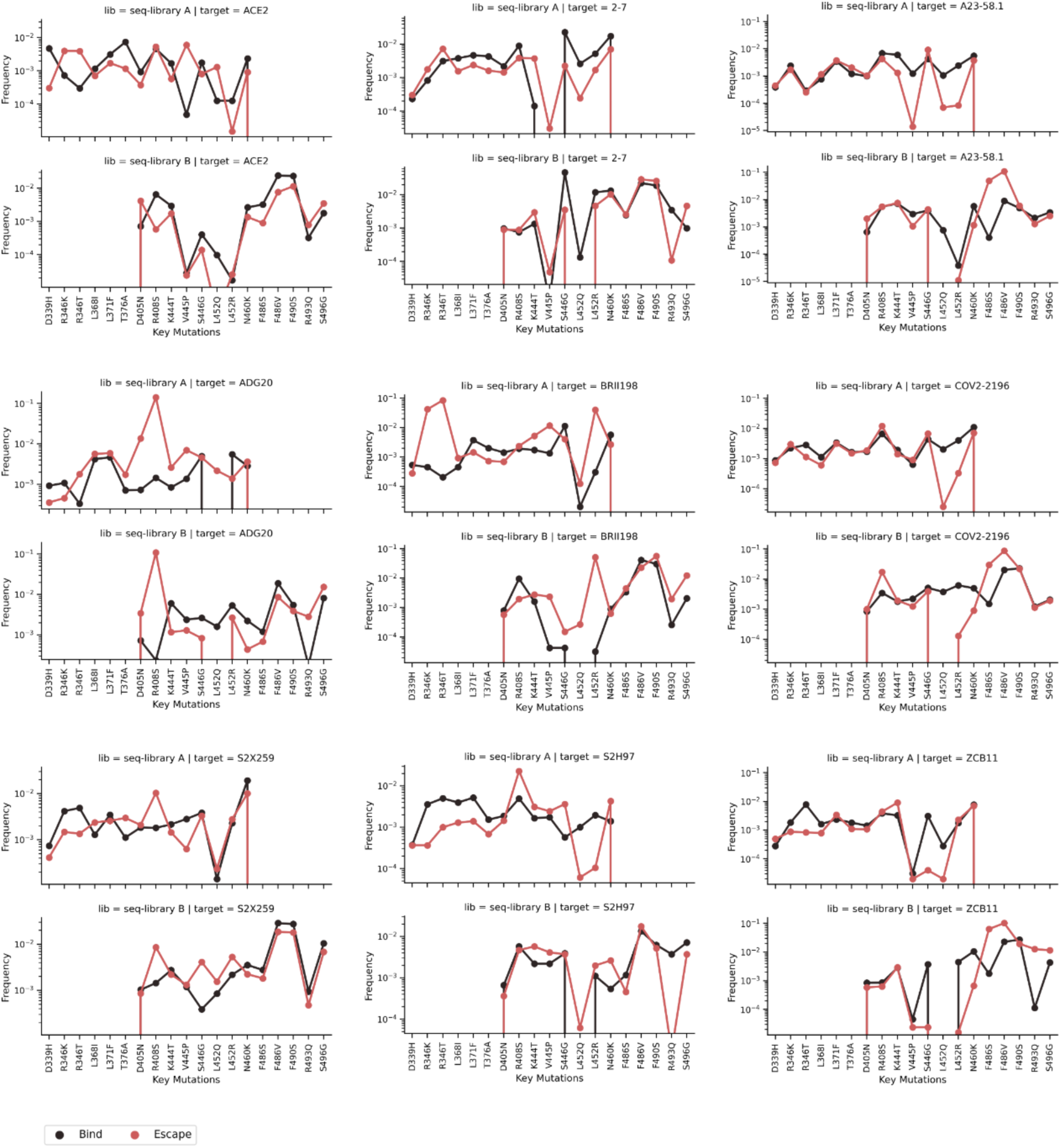
Line plots show the frequencies of selected mutations in the binding and escape fractions of the deep sequencing data. The selected mutations have been observed in previously identified Omicron sublineages.

**Supplementary Figure 5.**
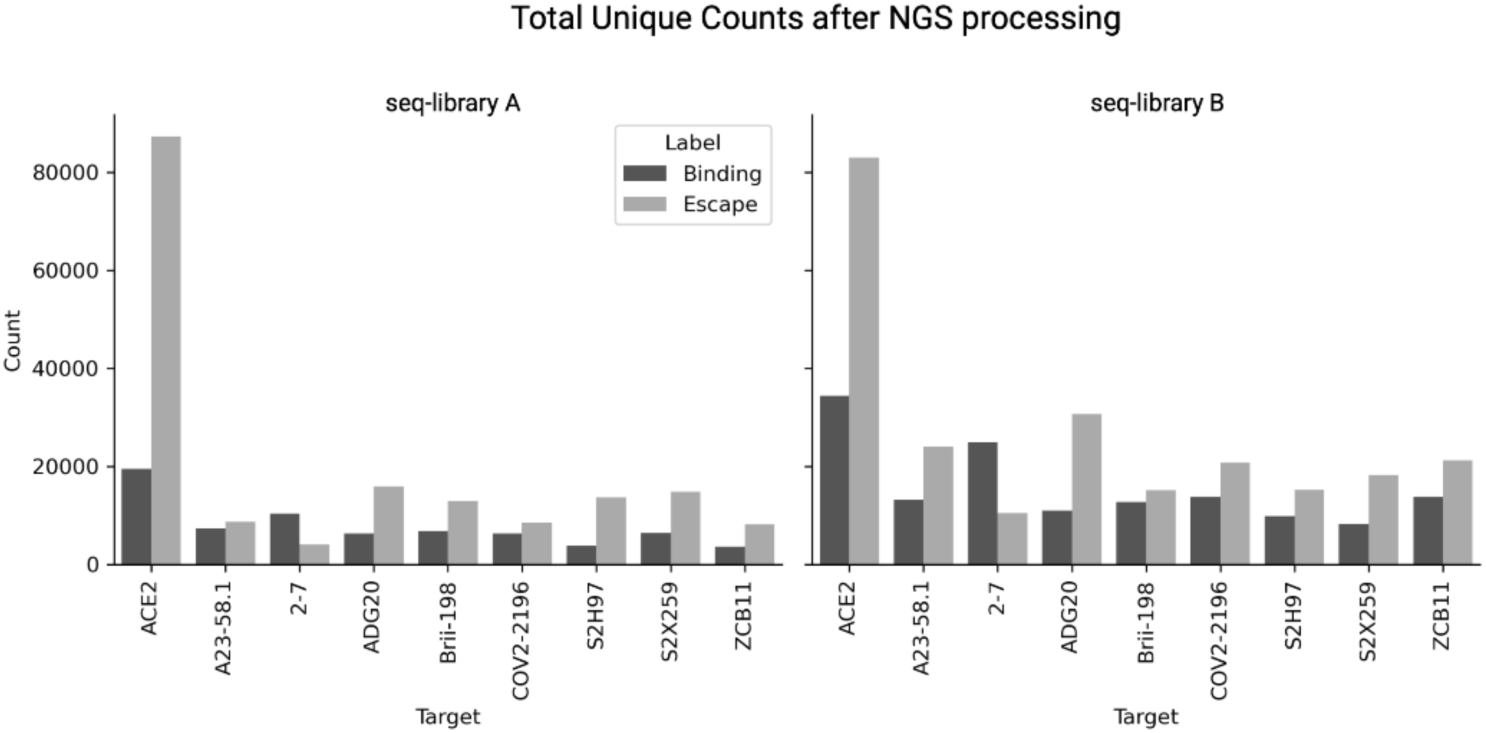
Total unique sequences (aa) in each deep sequencing dataset (following pre-processing).

**Supplementary Figure 6.**
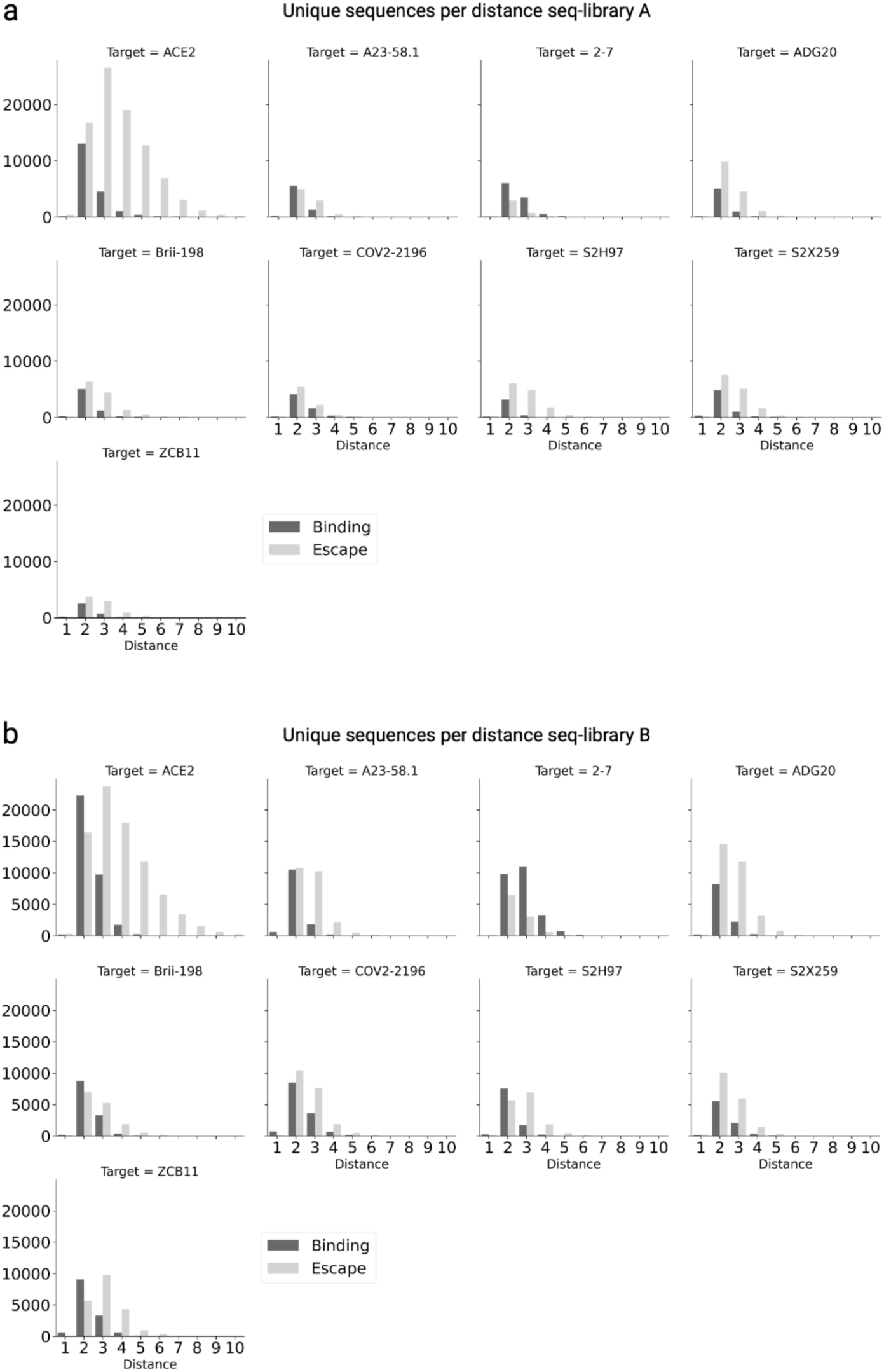
Number of unique sequences (aa) in each dataset per ED from WT BA.1 RBD sequence. To allow visual comparison between datasets, the maximum of the y-axis in all antibody datasets has been set to the highest count in all datasets (20,000).

**Supplementary Figure 7.**
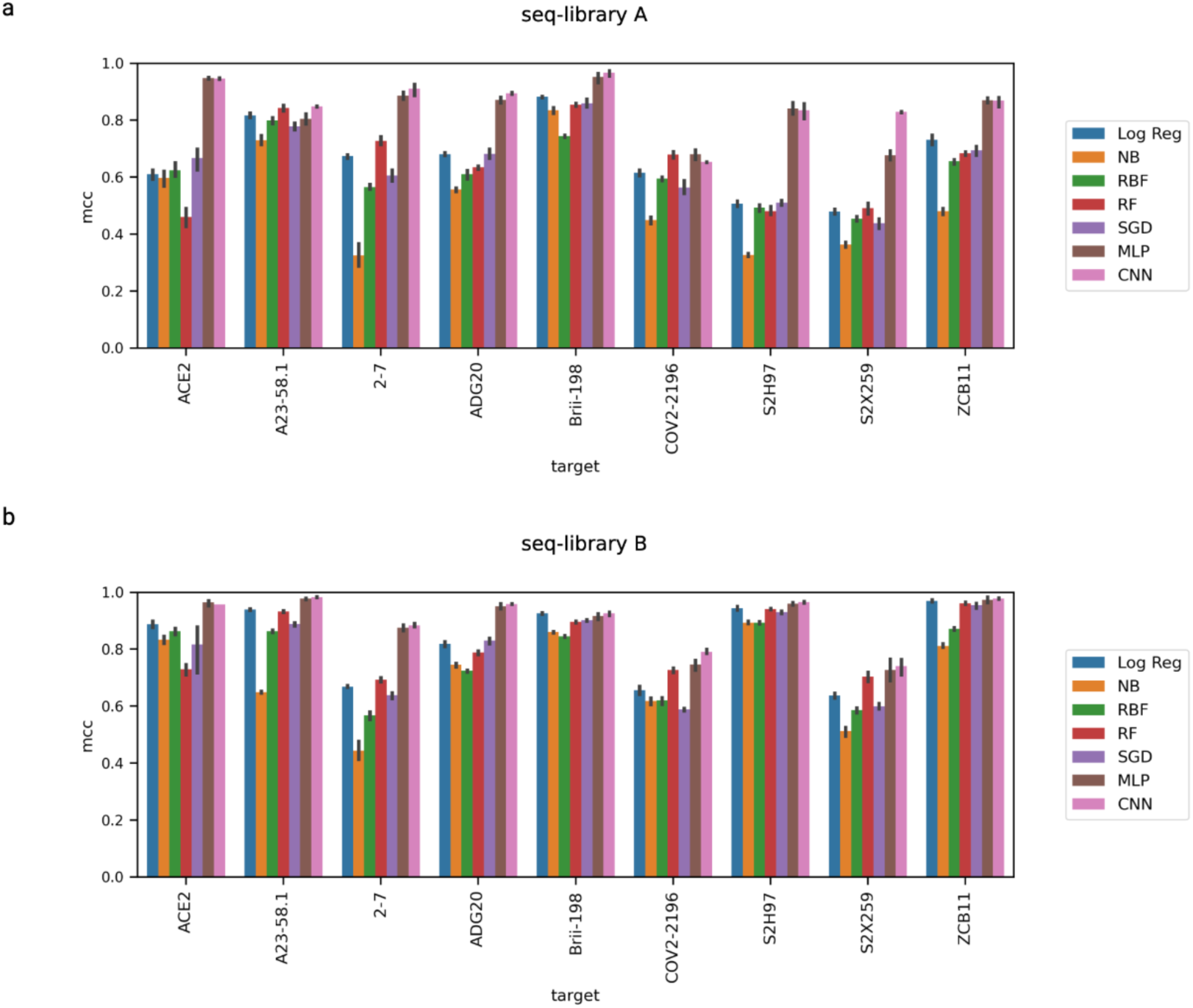
Barplots show MCC scores of all baseline machine learning models: Logistic Regression (Log Reg), Naive Bayes (NB), Radial Basis Function kernel SVM (RBF), Random Forest (RF), Stochastic Gradient Descent (SGD), and deep learning models: MLP and CNN, for **a**, seq-library A and **b**, seq-library B. All scores were evaluated through 5-fold cross-validation with a 80/10/10 train-val-test split.

**Supplementary Figure 8.**
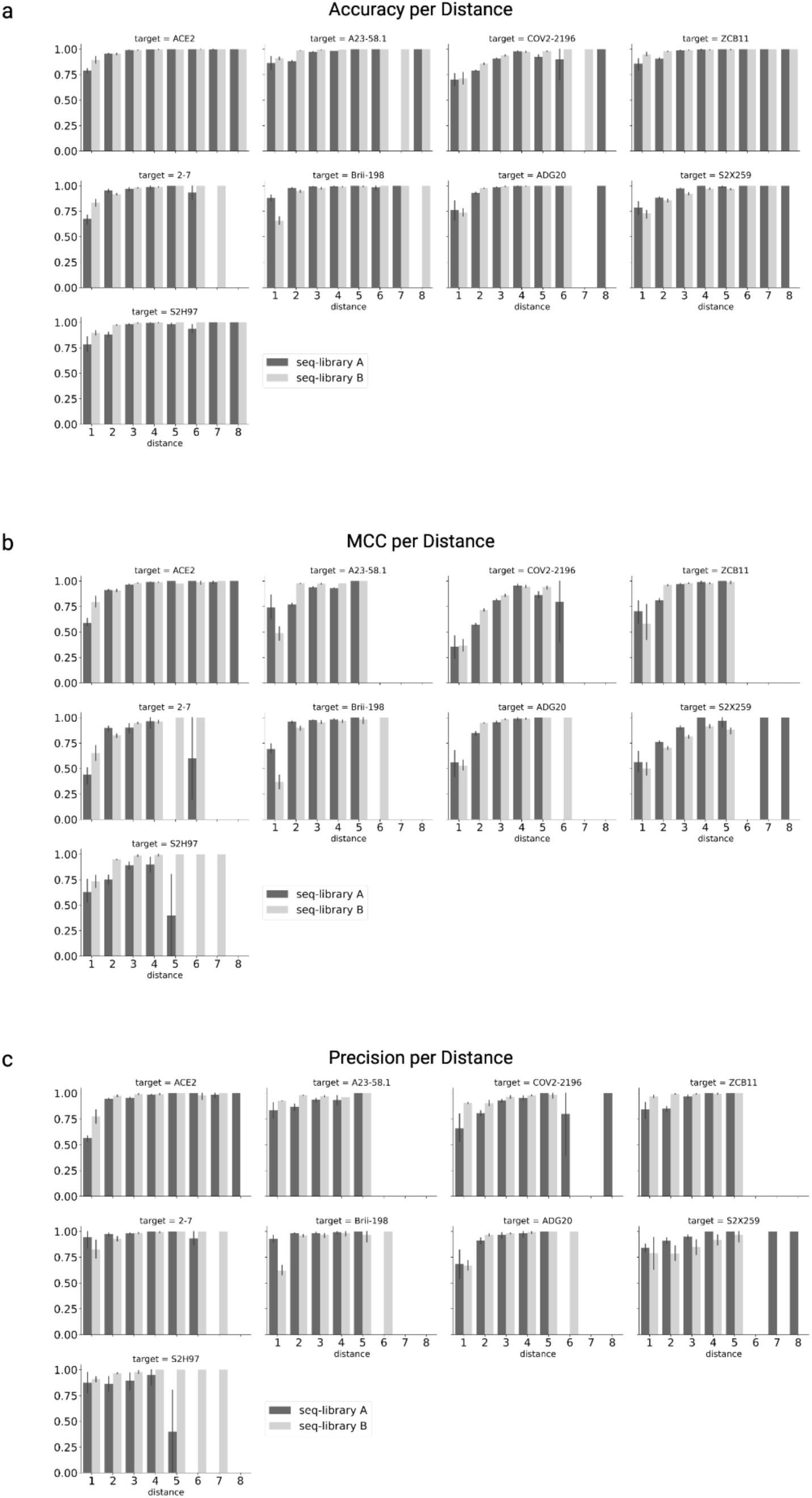
CNN model performances on test sequences based on ED from BA.1; shown are**a**, accuracy, **b**, MCC, and **c**, precision. All scores shown are combined results from 5-fold cross-validation with a 80/10/10 train-val-test split.

**Supplementary Figure 9.**
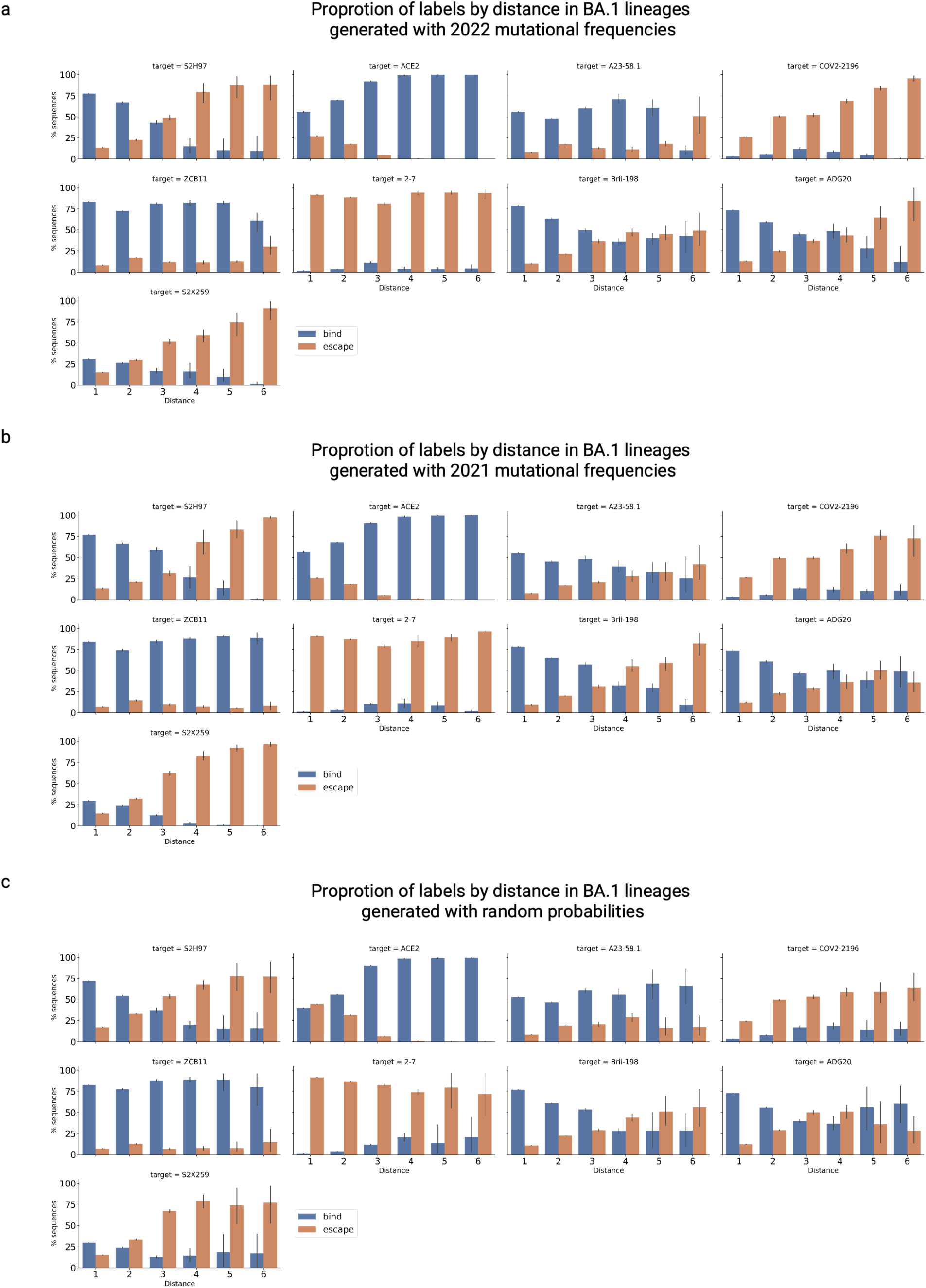
Percent of predicted binding and escape variants per ED (from BA.1) for each antibody. Predictions were run on 10 sets of synthetic lineages: BA.1-derived lineages based on GISAID mutational frequencies from **a**, 2021, **b**, 2022 or **c**, randomized probabilities (see Methods). Each synthetic lineage contains up to 250,000 sequences.

**Supplementary Figure 10.**
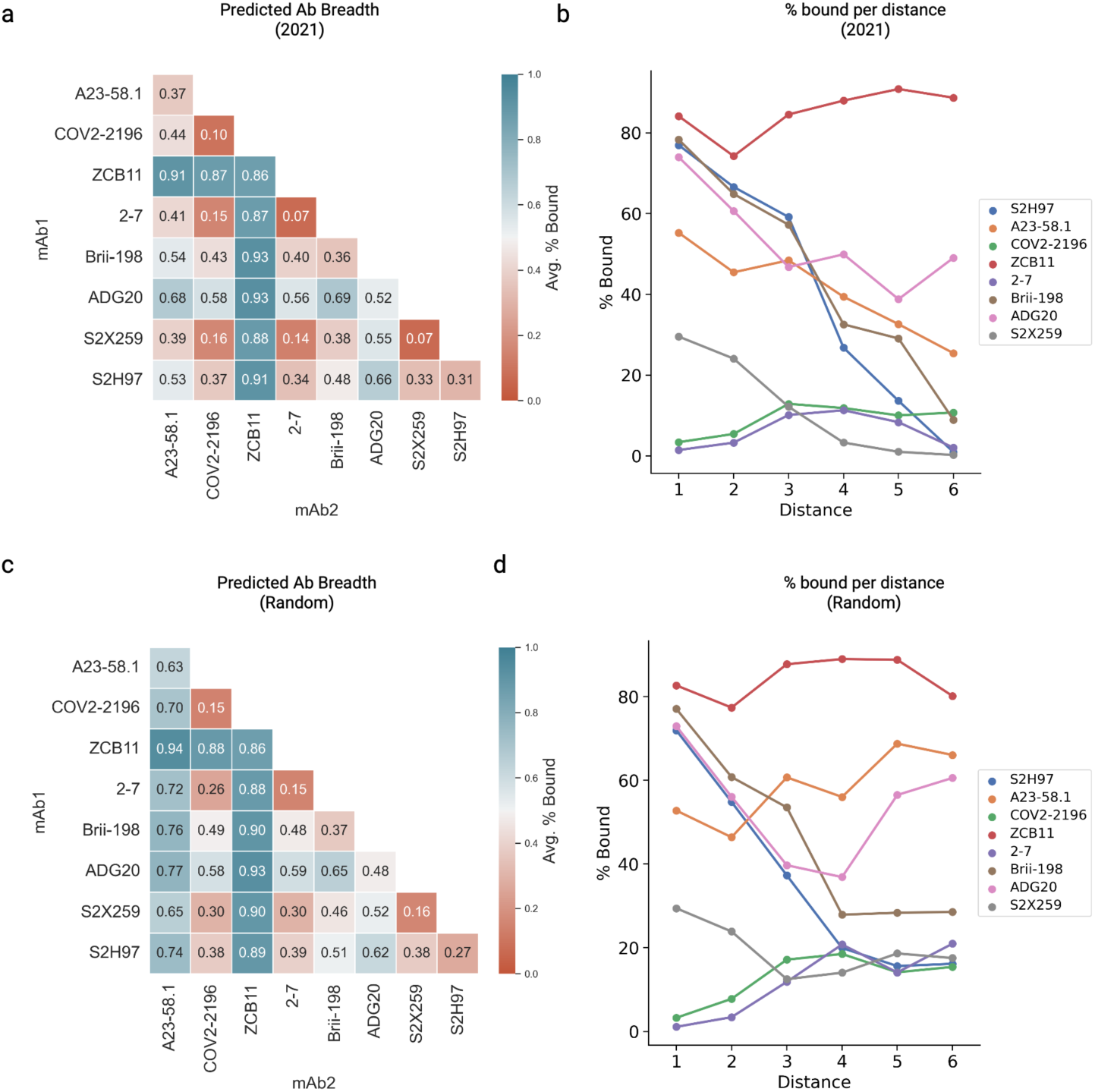
**a**, predicted total antibody breadth and **b**, antibody breadth per ED (from BA.1) on synthetic lineages (BA.1-derived lineages based on 2021 GISAID mutational frequencies, see Methods). **c**, predicted total antibody breadth and **d**, antibody breadth per ED (from BA.1) on randomized synthetic lineages (BA.1-derived lineages based on uniform random sampling frequencies, see Methods).

**Supplementary Figure 11.**
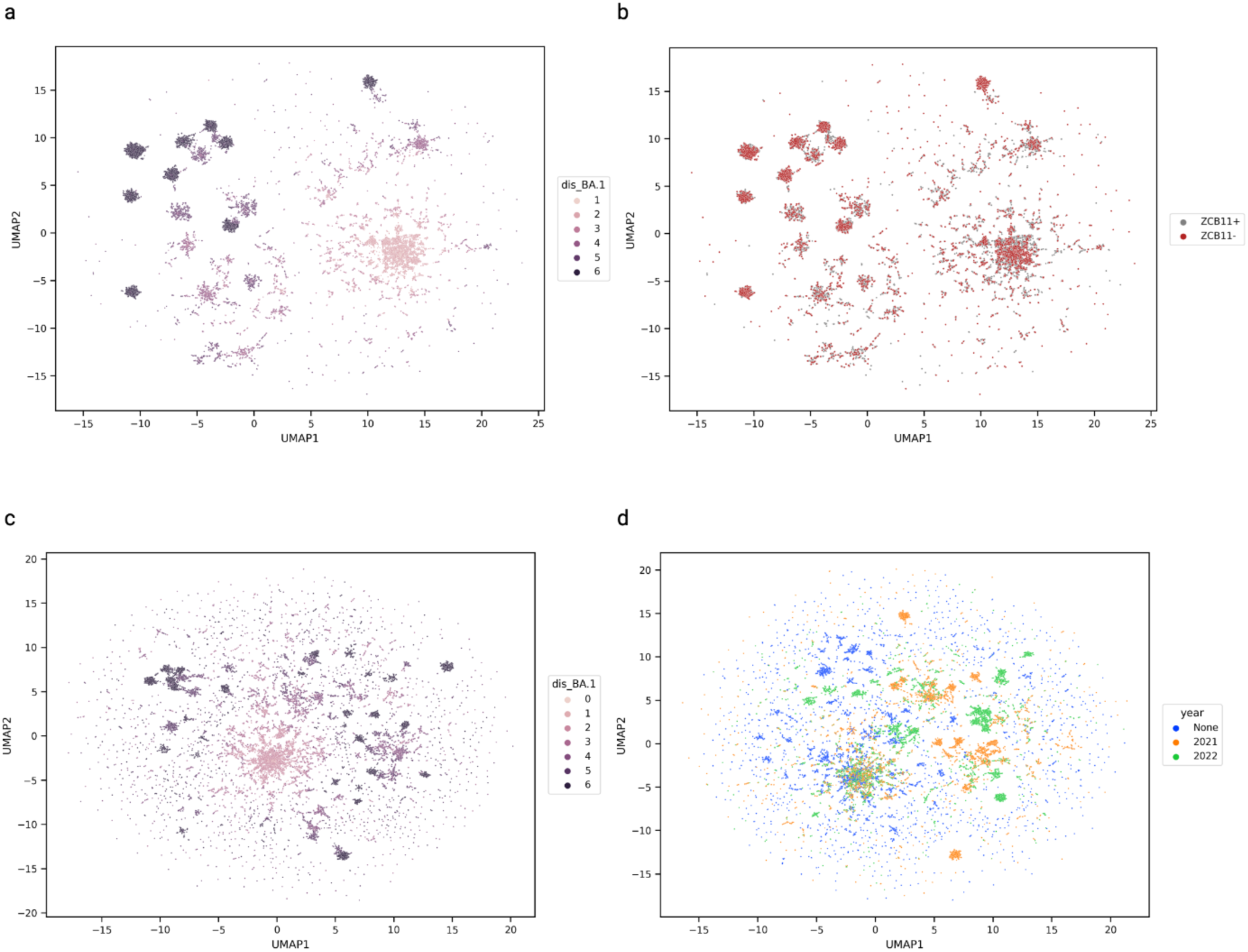
UMAPs show synthetic lineage variants in protein sequence space. **a**,sequences from Figure 4e, coloured to indicate their ED relative to WT (BA.1), and **b**, coloured to highlight sequences that bind (in gray) or escape (in red) from ZCB11. **c**, dimensionality reduced subsample of sequences taken from synthetic lineages from 2021, 2022 or random (none) probabilities coloured by their ED (from BA.1) and **d**, by the probabilities used to generate each lineage (“None” indicates sequences that were generated by random sampling from a uniform probability distribution across the RBD, with all aa substitutions allowed)

**Supplementary Figure 12.**
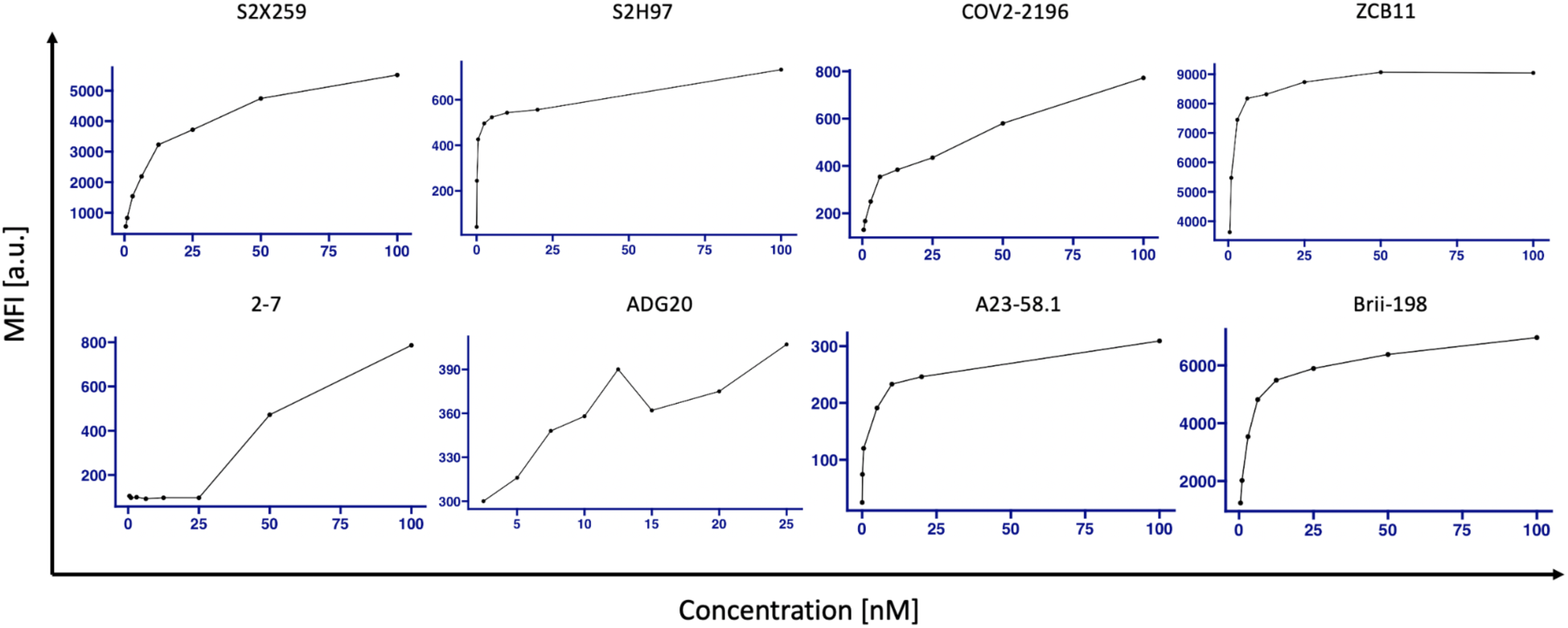
Titration curves of individual antibodies tested against yeast-displayed Omicron BA.1 RBD.

## SUPPLEMENTARY TABLES

**Supplementary Table 1.**
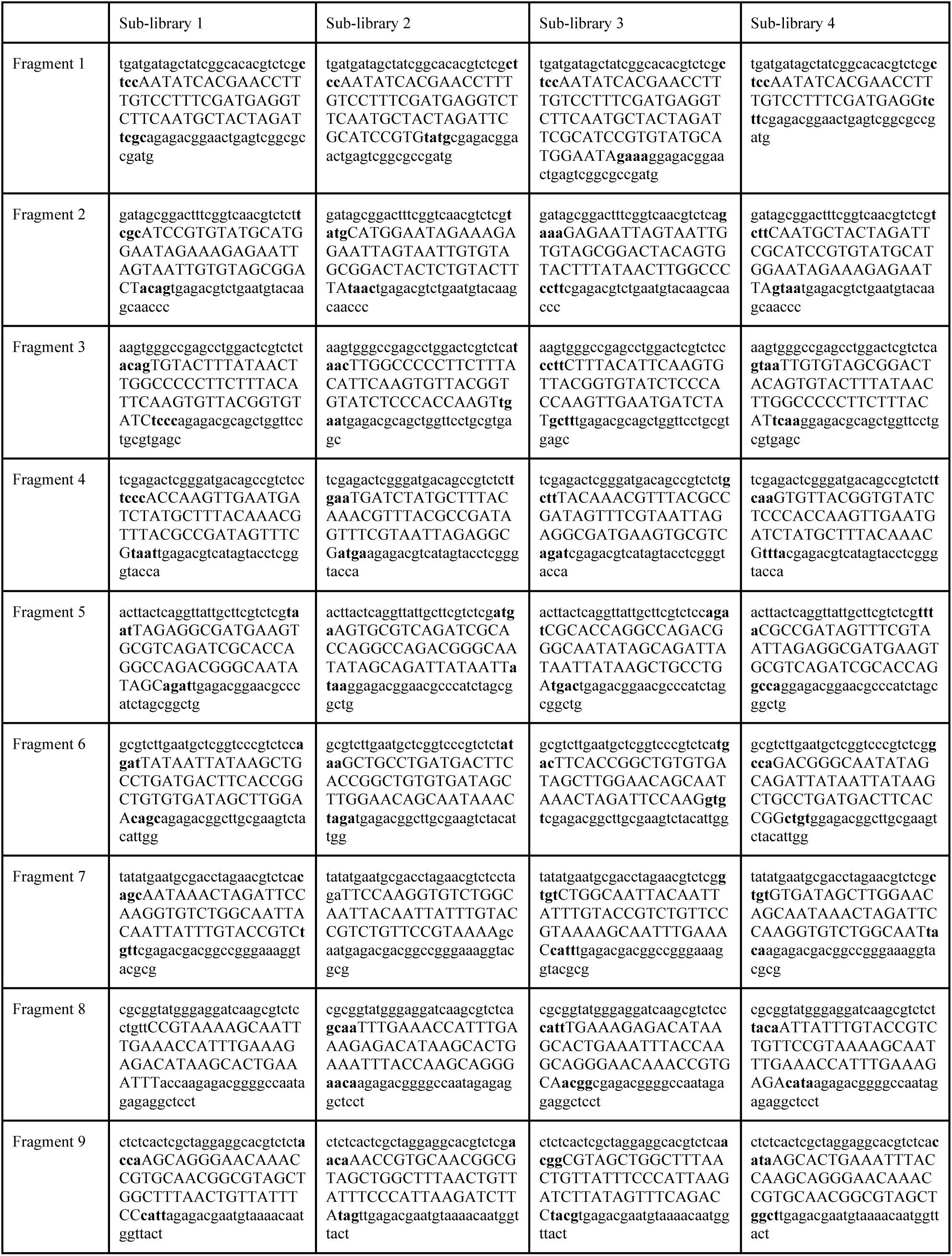
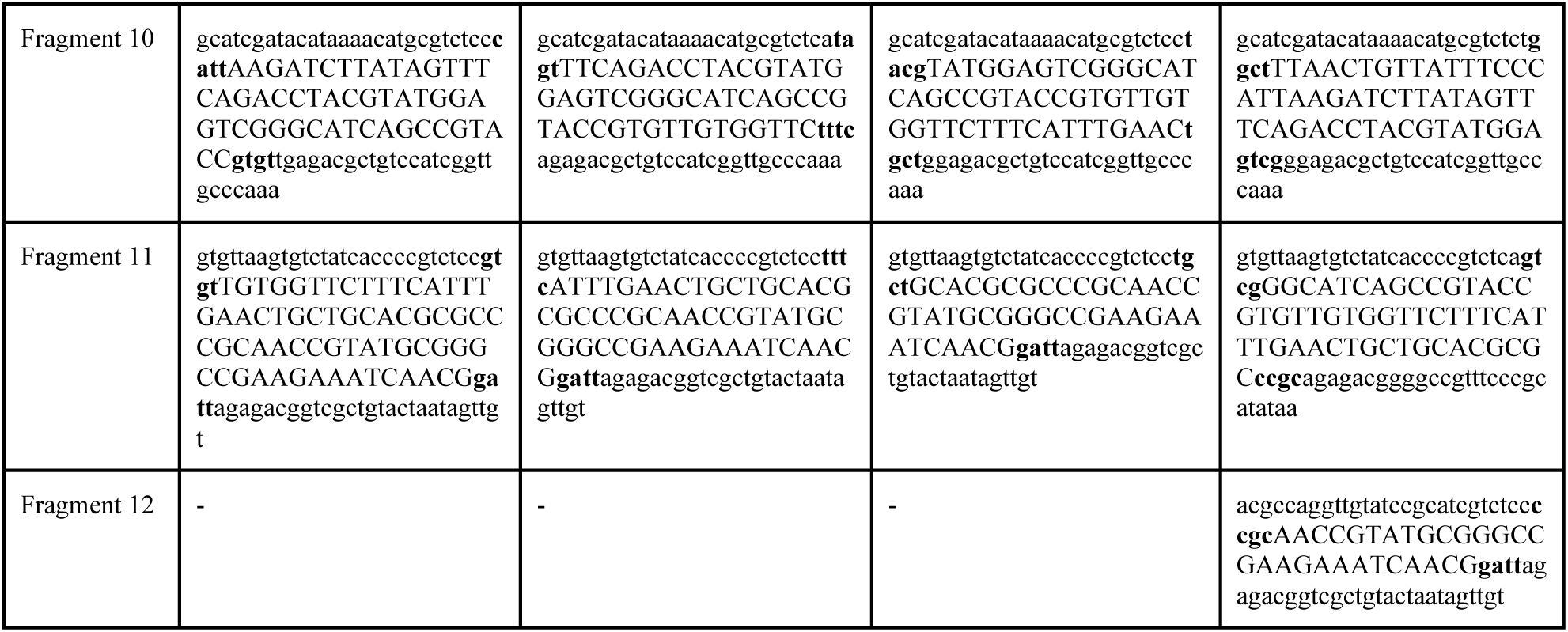
Sequences for fragments by sub-library. Sequences marked with uppercase letters are derived from the RBD open reading frame. The NNK codons are exclusively in this region. Bold lowercase sequences are the four nt homologies for GGA. The remaining lowercase sequences contain BsmBI recognition sites and primer binding sites for double strand synthesis.

**Supplementary Table 2:** Antibody concentrations used for FACS of yeast displayed RBD libraries. The concentrations were determined based on the titration curves shown in Supplementary Fig. 12.

| Therapeutic antibodies | Concentration [nM] |
| --- | --- |
| S2X259 | 12.5 |
| S2H97 | 2.5 |
| COV2-2196 | 12.5 |
| ZCB11 | 6.25 |
| 2-7 | 60 |
| ADG20 | 7.5 |
| A23-58.1 | 5 |
| Brii-198 | 10 |

**Supplementary Table 3:** Deep sequencing statistics for sorted RBD libraries.

| Population | Antigen | Paired and filtered Reads |  |
| --- | --- | --- | --- |
|  |  | Binding | Non-Binding |
| seq-Library A | ACE2 | 1.92E+06 | 1.75E+06 |
| seq-Library B | ACE2 | 2.32E+06 | 1.60E+06 |
| seq-Library A, ACE2 binding | 2-7 | 3.27E+05 | 5.52E+05 |
| seq-Library A, ACE2 binding | A23-58.1 | 1.20E+06 | 1.08E+06 |
| seq-Library A, ACE2 binding | ADG20 | 8.37E+05 | 8.83E+05 |
| seq-Library A, ACE2 binding | Brii-198 | 9.40E+05 | 3.59E+05 |
| seq-Library A, ACE2 binding | COV2-2196 | 5.59E+05 | 9.40E+05 |
| seq-Library A, ACE2 binding | S2H97 | 5.37E+05 | 7.23E+05 |
| seq-Library A, ACE2 binding | S2X259 | 6.22E+05 | 5.59E+05 |
| seq-Library A, ACE2 binding | ZCB11 | 5.52E+05 | 8.30E+05 |
| seq-Library B, ACE2 binding | 2-7 | 1.02E+06 | 9.32E+05 |
| seq-Library B, ACE2 binding | A23-58.1 | 9.12E+05 | 7.38E+05 |
| seq-Library B, ACE2 binding | ADG20 | 9.38E+05 | 9.03E+05 |
| seq-Library B, ACE2 binding | Brii-198 | 9.70E+05 | 5.48E+05 |
| seq-Library B, ACE2 binding | COV2-2196 | 7.66E+05 | 9.70E+05 |
| seq-Library B, ACE2 binding | S2H97 | 8.42E+05 | 4.43E+05 |
| seq-Library B, ACE2 binding | S2X259 | 7.50E+05 | 7.66E+05 |
| seq-Library B, ACE2 binding | ZCB11 | 9.32E+05 | 6.09E+05 |

**Supplementary Table 4:** Hyperparameter Search Conditions for CNN and MLP models.

| Parameter | MLP | ProtCNN |
| --- | --- | --- |
| Learning Rate | 0.01 - 0.0001 |  |
| Optimizer | Adam, SGD |  |
| Minority Ratio (dataset balance) | 0.1 - 0.5 |  |
| Epochs | 15 - 75 |  |
| Test/Val ratio | 0.1-0.25 |  |

| MLP Parameters |  |
| --- | --- |
| Dense Dimensions | 32 - 512 |
| # Dense Layers | 1-3 |
| Dense Dropout | 0 - 0.5 |

| CNN Parameters |  |  |
| --- | --- | --- |
| Kernel Size |  | 3 - 21 |
| Stride |  | 1-3 |
| Filter Number |  | 32 - 512 |
| Padding |  | "Same", 1 |
| Pool Size |  | 1-3 |
| Pool Stride |  | 1-3 |
| Dilation Rate |  | 2-5 |
| Residual Blocks |  | 1-3 |

**Supplementary Table 5:**
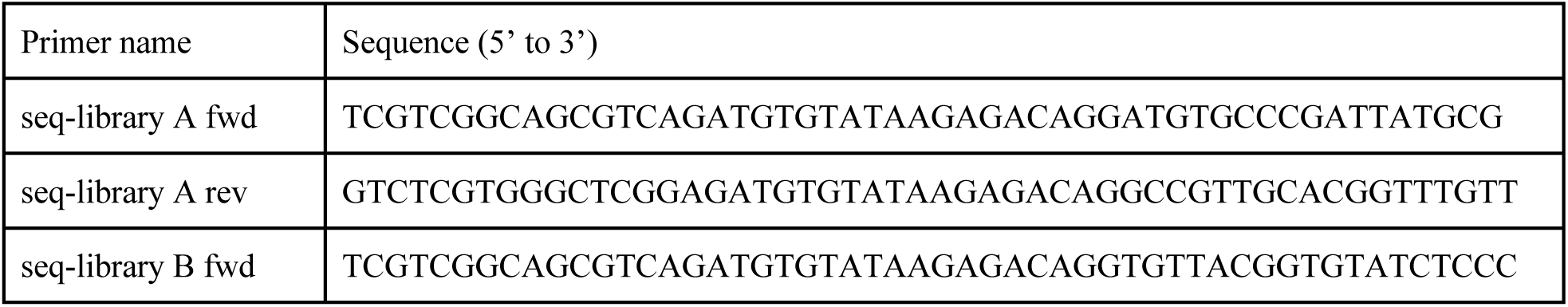
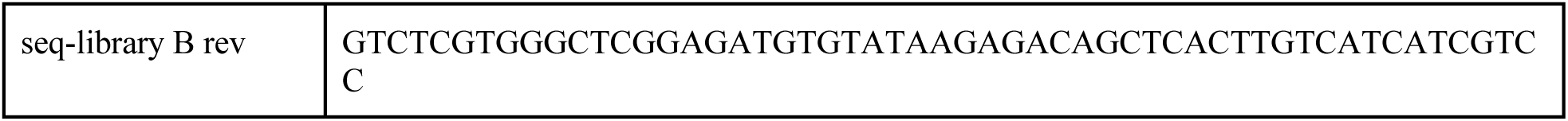
Primers used to amplify seq-libraries A and B in a targeted fashion for subsequent deep sequencing.

## Supplementary File

Experimentally measured binding affinity and neutralization of antibodies against SARS-CoV-2 variants from publications.

